# Loss of *NF1* causes tactile hypersensitivity and impaired synaptic transmission in a *Drosophila* model of autism spectrum disorder

**DOI:** 10.1101/2022.03.04.482984

**Authors:** Alex Dyson, Shruti Garg, D. Gareth Evans, Richard A. Baines

## Abstract

Autism Spectrum Disorder (ASD) is a neurodevelopmental condition in which the mechanisms underlying its core symptomatology are largely unknown. Studying animal models of monogenic syndromes associated with ASD, such as neurofibromatosis type 1 (NF1), can offer insights into its aetiology. Here, we show that loss of function of the *Drosophila NF1* ortholog results in larval tactile hypersensitivity, paralleling the sensory abnormalities observed in individuals with ASD. Mutant larvae also exhibit synaptic transmission deficits at the glutamatergic neuromuscular junction (NMJ), with increased spontaneous but reduced evoked release. Diminished expression of *NF1* specifically within central cholinergic neurons induces both excessive neuronal firing and tactile hypersensitivity, suggesting the two may be linked. Furthermore, both impaired synaptic transmission and behavioural deficits are fully rescued via knockdown of Ras proteins. These findings validate *NF1*^*-/-*^ *Drosophila* as a tractable model of ASD with the potential to elucidate important pathophysiological mechanisms.

## Introduction

Autism spectrum disorder (ASD) is a neurodevelopmental disorder characterised by social communication deficits and repetitive behaviours (American Psychiatric Association, 2013), affecting 1 – 2% of the population (Baio et al., 2018). The condition incurs a substantial impact on quality of life (Brugha et al., 2011; Fortuna et al., 2016; Howlin and Moss, 2012), and carries a considerable economic burden (Knapp et al., 2009). However, there are currently no effective treatments that target the core behavioural symptoms due to a poor understanding of the underlying pathological mechanisms (Vorstman et al., 2014).

While the majority (∼75%) of ASD cases are idiopathic, approximately 4% occur in association with a monogenic neurodevelopmental syndrome (Miles et al., 2005; Tammimies et al., 2015). Because monogenic forms of ASD have a known single causative mutation, they are comparatively simple to model and are thus highly tractable from a biomedical research perspective (Sztainberg and Zoghbi, 2016). One such condition is Neurofibromatosis Type 1 (NF1), an autosomal dominant disorder arising from loss of function mutations in the *NF1* gene on chromosome 17 (Gutmann et al., 2017). Although typically classified as a cancer predisposition syndrome, cognitive and behavioural issues are common in NF1 (Hyman et al., 2005), and the prevalence of ASD amongst affected individuals is estimated at 25% (Garg et al., 2013). Furthermore, the symptomatology of ASD amongst the NF1 population is highly similar to that in idiopathic individuals, as is the degree of symptom heterogeneity (Morris et al., 2016).

*NF1* encodes for neurofibromin, a ∼320 kDa multi-domain cytoplasmic protein expressed most strongly in oligodendrocytes, Schwann cells, and neurons (Daston et al., 1992; Marchuk et al., 1991). While its best characterised molecular function is as a Ras-GTPase Activating Protein (Ras-GAP), neurofibromin also functions in numerous additional pathways including as an activator of cAMP/PKA signalling (Ratner and Miller, 2015). At the cellular level, *NF1* is required for neural connectivity (Shofty et al., 2019), dendritic spine formation (Shih et al., 2020), cell migration (Sanchez-Ortiz et al., 2014), and synaptic transmission (Costa et al., 2002; Cui et al., 2008; Molosh et al., 2014; Omrani et al., 2015; Shilyansky et al., 2010), all processes which may be disrupted in ASD (Yenkoyan et al., 2017). Several studies in mouse models of NF1 have studied how altered neurotransmission in the CNS results in cognitive and behavioural impairments. Typically, these postulate an increase in GABAergic transmission and a consequent deficit in long-term potentiation (Costa et al., 2002; Cui et al., 2008; Molosh et al., 2014; Omrani et al., 2015). These changes involve both increased ERK-mediated phosphorylation of synapsin (Cui et al., 2008) and a reduction in neuronal I_h_ current (Omrani et al., 2015).

In contrast, how *NF1* regulates excitatory transmission, and the impact of this on behaviour, is less clear. Given that ASD-associated genes appear to be predominantly expressed in glutamatergic neurons during development (Parikshak et al., 2013; Willsey et al., 2013), resolving this question may shed light on how aberrant synaptic transmission underlies core ASD symptomatology. Here, we identify an ASD-relevant behavioural phenotype in *NF1*^*P1*^ *Drosophila melanogaster* larvae, namely an increased likelihood of exhibiting a nocifensive response when exposed to a typically innocuous mechanical stimulus (tactile hypersenstivity). Furthermore, we show that, at the glutamatergic larval neuromuscular junction (NMJ), spontaneous transmitter release is increased while evoked release is reduced. While the latter is homeostatically compensated for by a postsynaptic increase in muscle input resistance (R_in_), the former may reflect neuronal hyperexcitability. Indeed, in semi-intact larval preparations, in which the central nervous system (CNS) is intact, we observe excessive endogenous activity at the NMJ. Both neuronal hyperexcitability and tactile hypersensitivity arise due to loss of *NF1* in the CNS, rather than in peripheral or sensory neurons. Lastly, we show that knockdown of either *Ras85D* or *Ras64B* in *NF1*^*P1*^ larvae is sufficient to fully rescue these phenotypes, indicating that they arise from excessive Ras/MAPK pathway signalling.

## Results

### *NF1*^*P1*^ larvae exhibit hypersensitivity to mechanical stimulation

Altered sensory sensitivity is common in ASD and included in the DSM-V criteria under the domain of repetitive behaviours and restricted interests (American Psychiatric Association, 2013). Therefore, we investigated whether *NF1* mutant *Drosophila* larvae exhibit altered responses to stimuli relative to isogenic K33 controls. Following exposure to noxious stimuli such as heat or mechanical touch, wildtype third instar larvae display a stereotypic, ‘corkscrew-like’ rolling behaviour that is suggested to be a nocifensive response evolved to protect the animal from parasitic wasp attack (Figure 1A) (Hwang et al., 2007; Tracey et al., 2003). Assaying this behaviour has led to the identification of several genes required for appropriate sensory transduction and processing (Mauthner et al., 2014; Tracey *et al*., 2003; Walcott et al., 2018). When a brief mechanical stimulation (see Methods; Figure 1A) is applied to the posterior end of a K33 larva, this generally does not elicit a response. In contrast, the number of *NF1*^*P1*^ larvae exhibiting a nocifensive response is significantly greater (Figure 1B). Typical responses for K33 and *NF1*^*P1*^ larvae are shown in Videos S1 and S2, respectively. A similar phenotype was seen when comparing the *NF1*^*E2*^ line to its *w*^*1118*^ isogenic control (Figure 1C), and pan-neuronal overexpression of *NF1* fully rescues this phenotype (Figure 1D).

**Figure 1.**
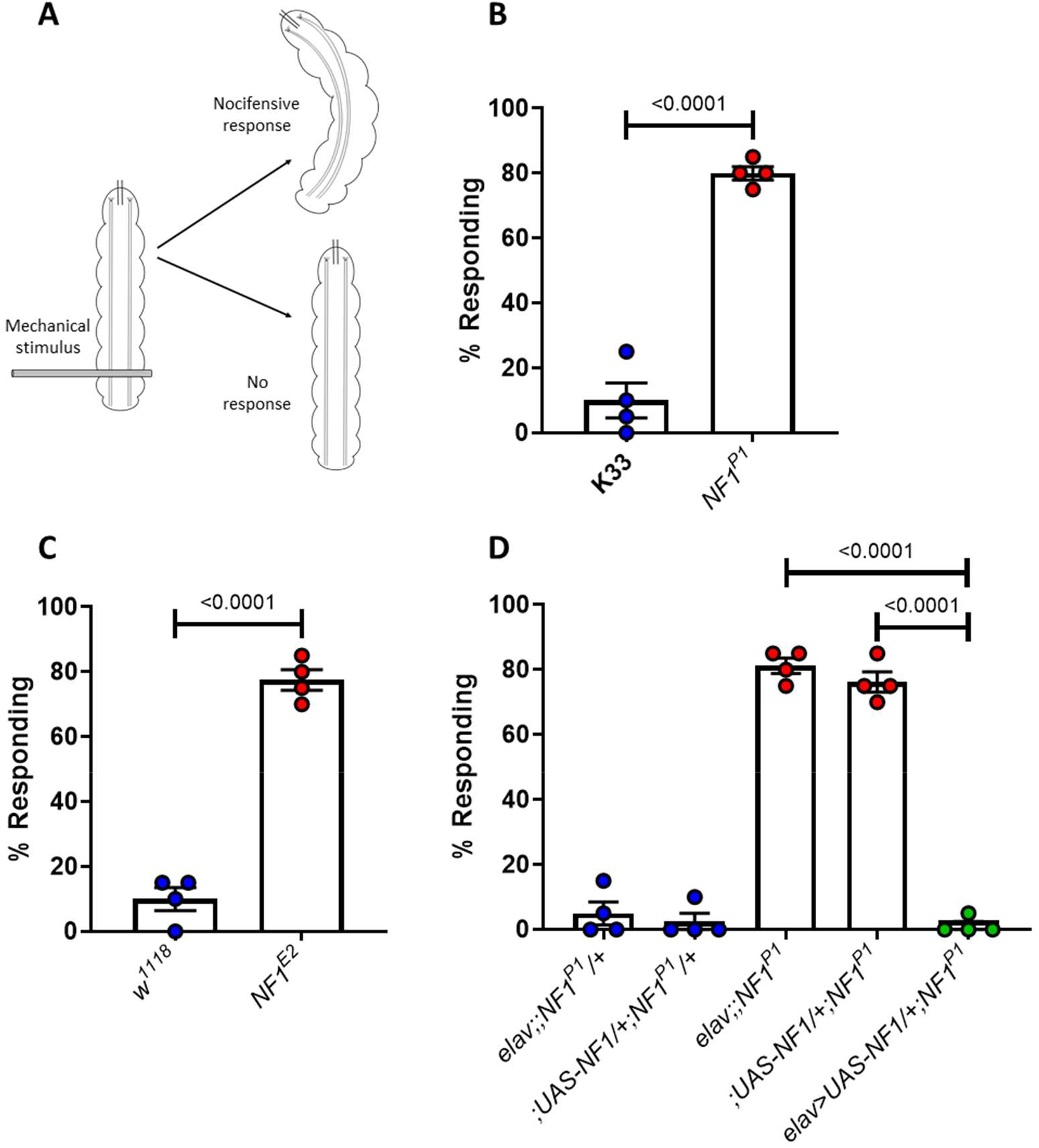
*NF1*^*-/-*^ larvae display hypersensitivity to mechanical stimulation. *A)* Schematic of the mechanoreception assay used to characterise tactile hypersensitivity. An insect pin is pressed down firmly across the posterior end of a larva, an action that may or may not induce a nocifensive rolling motion. *B)* The mean percentage of larvae (*n* = 4 trials, 20 larvae per trial) responding to mechanical stimulation is significantly greater in *NF1*^*P1*^ than in the K33 control. *C)* A similar effect is seen when comparing *NF1*^*E2*^ to *w*^*1118*^ controls. *D)* Pan-neuronal overexpression of *UAS-NF1* in the *NF1*^*P1*^ background fully rescues the phenotype. All data are presented as mean ± SEM. Statistical comparisons were made via either an unpaired, two-tailed student’s t-test (panels B and C) or a one-way ANOVA followed by Tukey’s post-hoc test (panel D).

### *NF1*^*P1*^ larvae exhibit increased spontaneous but reduced evoked transmission at the NMJ

Given that *NF1* regulates an ASD-relevant behavioural phenotype in *Drosophila* larvae, and defects in synaptic transmission are thought to contribute to ASD symptomatology (Zoghbi and Bear, 2012), we next investigated whether synaptic transmission was altered. To this end, we made use of the *Drosophila* larval NMJ. This is an easily-accessible and well-characterised model glutamatergic synapse at which the proteins governing synaptic transmission exhibit a very high degree of conservation with their mammalian counterparts and function in a manner highly similar to those at mammalian central synapses (Keshishian et al., 1996). Furthermore, defects in synaptic transmission at the NMJ have been observed following the mutation of *Drosophila* orthologs of other human ASD-susceptibility genes (Russo and DiAntonio, 2019; Tsai *et al*., 2012; Valdez et al., 2015). Thus, while the NMJ is a peripheral synapse, in *Drosophila* it provides an excellent system with which to study ASD-relevant changes in excitatory transmission.

Previous work (Tsai et al., 2012) characterised altered synaptic transmission at the NMJ of *NF1*-null larvae in a reduced CaCl_2_ concentration. To more accurately reflect the physiological environment, we first carried out current-clamp recordings in HL3 saline containing 1.5 mM CaCl_2_ (Stewart et al., 1994). Under such conditions, no significant difference in evoked excitatory junction potential (EJP) amplitude was observed between *NF1*^*P1*^ and K33 lines. However, in *NF1*^*P1*^ larvae, both the frequency and amplitude of miniature EJPs (mEJPs; i.e. membrane depolarisations in response to the spontaneous release of transmitter) were significantly increased, while quantal content (the number of vesicles released by an action potential) was significantly reduced (Figure 2A-F). The same phenotypes were also observed in *NF1*^*E2*^ larvae when compared to their isogenic control line *w*^*1118*^ (Figure 2G-L).

**Figure 2.**
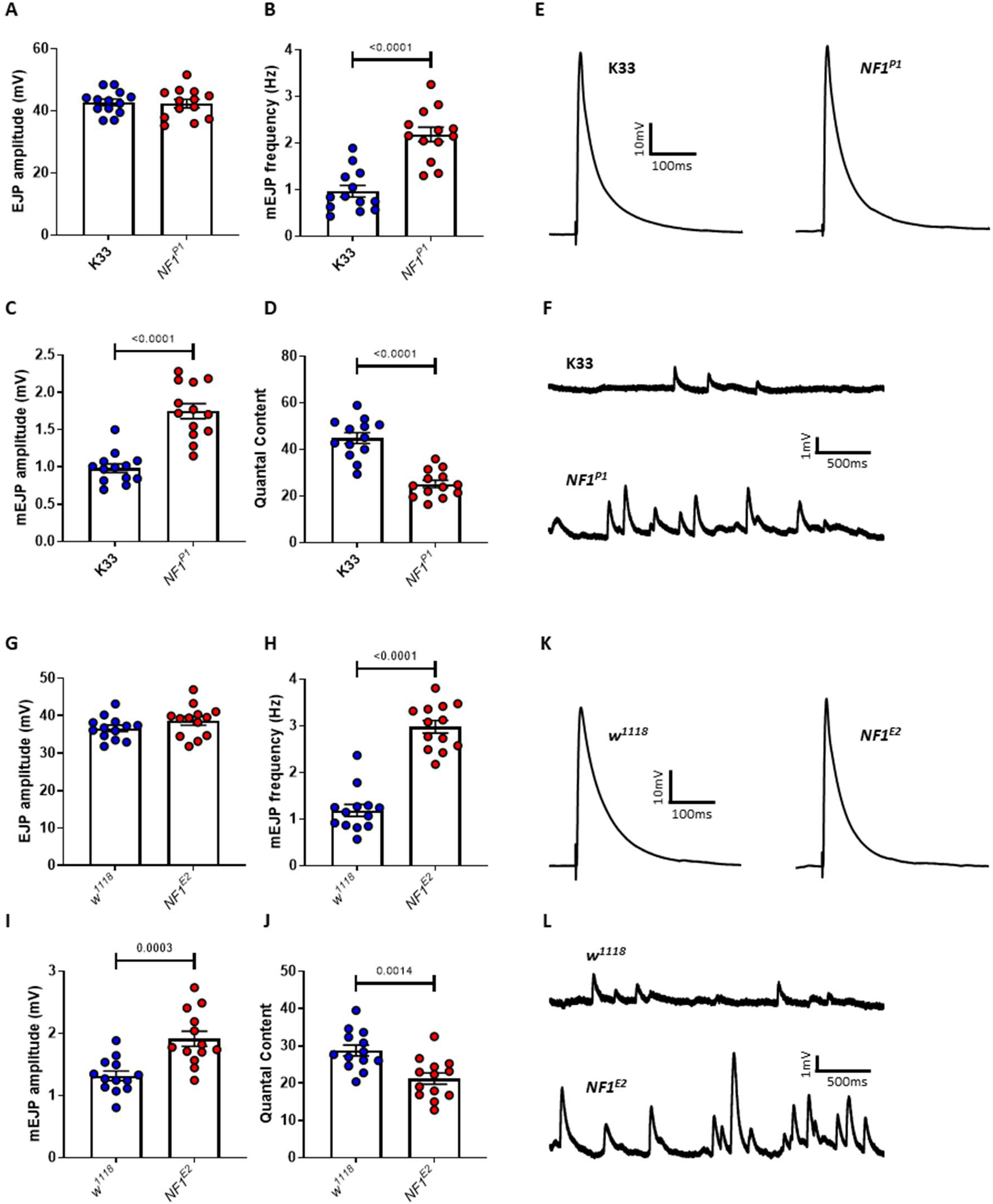
*NF1*^*-/-*^ mutants display reduced evoked but increased spontaneous excitatory synaptic transmission. *A)* Under current-clamp, EJP amplitude is not significantly altered (p=0.82) in *NF1*^*P1*^ mutants compared to that of K33 controls. However, both *B)* mEJP frequency and *C)* mEJP amplitude are significantly increased, while *D)* quantal content is significantly decreased. *E-F)* Representative traces of EJPs and mEJPs, respectively, for K33 and *NF1*^*P1*^ lines. *G–J)* A similar phenotype is seen in *NF1*^*E2*^ mutants compared to *w*^*1118*^ controls, with no change in EJP amplitude (p=0.18), a significant increase in mEJP frequency and mEJP amplitude, and a significant decrease in quantal content. *K–L)* Representative traces of EJPs and mEJPs, respectively, analysed in G-J. All data are presented as mean ± SEM. All statistical comparisons were made via unpaired, two-tailed student’s *t-*test.

Increased mEJP frequency and amplitude were recapitulated by pan-neuronal (*elav>NF1*^*RNAi*^*;Dicer2*), but not muscular (*MHC>NF1*^*RNAi*^*;Dicer2*), knockdown of *NF1* (Figure 3A-F). We did observe a slight but significant decrease in EJP amplitude in *elav>NF1*^*RNAi*^*;Dicer2* larvae, as well as a significant decrease in quantal content in *MHC>NF1*^*RNAi*^*;Dicer2* larvae, though this was much smaller in magnitude than that seen following either presynaptic *NF1* knockdown or in homozygous *NF1* deletion mutants (i.e. *NF1*^*P1*^ or *NF1*^*E2*^ mutants, c.f. Figure 2). Pan-neuronal knockdown of a second, independent, *UAS-NF1*^*RNAi*^ transgene showed a similar phenotype (Figure S1). Lastly, synaptic dysfunction was fully rescued *via* pan-neuronal overexpression of a *UAS-NF1* transgene in the *NF1*^*P1*^ background (Figure 3G-L). Together, these data strongly indicate that the observed transmission defects result from presynaptic loss of *NF1* activity, although a post-synaptic role for *NF1* in synaptic transmission cannot be ruled out.

**Figure 3.**
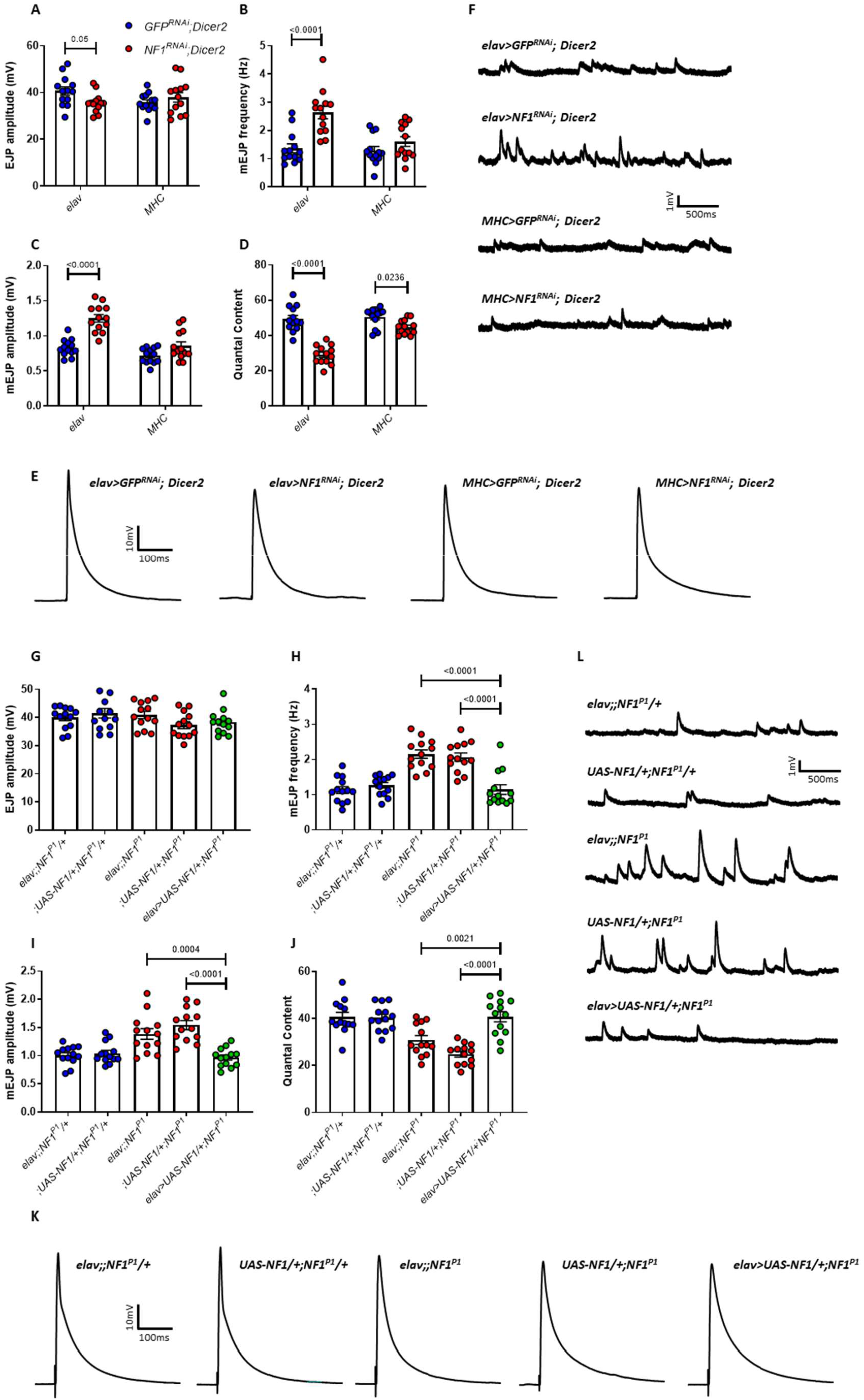
Synaptic transmission deficits are presynaptic in origin and specific to loss of *NF1* expression. *A)* EJP amplitude is significantly reduced following presynaptic knockdown (*elav*) of *NF1* via expression of *NF1*^*RNAi*^*;Dicer2*, whereas postsynaptic knockdown (*MHC*) has no effect (p=0.57). *B)* Similar to *NF1*^*P1*^ and *NF1*^*E2*^, mEJP frequency is significantly reduced following presynaptic but not postsynaptic (p=0.37) knockdown. *C)* Presynaptic *NF1* knockdown increases mEJP amplitude, whereas postsynaptic knockdown does not (p=0.063). *D)* Quantal content is significantly reduced following *NF1* knockdown both presynaptically and postsynaptically, although to a lesser extent in the latter. *E-F)* Representative traces of EJPs and mEJPs, respectively, analysed in *A-D. G-L)* Pan-neuronal overexpression of *UAS-NF1* via *elav-GAL4* in the *NF1*^*P1*^ background rescues synaptic transmission deficits, with no significant differences between the rescue line (green circles; *elav>UAS-NF1;NFI*^*P1*^) and either heterozygous control (blue circles) for any parameter examined. Furthermore, in panels H-J, both homozygous mutant controls (red circles) were significantly different to both heterozygous controls, and there were no significant differences between either of the heterozygous controls or either of the homozygous mutant controls, respectively. All data are presented as mean ± SEM. All statistical comparisons in A-D were made via two-way ANOVA followed by Sidak’s multiple comparisons test, in order to compare *NF1*^*RNAi*^*;Dicer2* and *GFP*^*RNAi*^*;Dicer2* larvae within each driver group. All comparisons in G-J were made via one-way ANOVA followed by Tukey’s multiple comparisons test.

In addition to recording in current-clamp mode, immediately following each recording in Figure 2A-F we recorded excitatory junction currents (EJCs) and miniature EJCs (mEJCs) from the same muscle by switching to voltage clamp (Figure 4A-F). Whilst, under current clamp, EJP amplitude is unchanged (Figure 2A), EJC amplitude was significantly reduced in *NF1*^*P1*^ larvae. By contrast, mEJC amplitude, increased under current clamp, was unchanged relative to K33 controls in voltage clamp. Consistently, however, mEJC frequency remained significantly increased and quantal content significantly decreased under both recording modes. Given that both sets of recordings were carried out from the same muscles, the discrepancy between (m)EJP and (m)EJC amplitudes in *NF1*^*P1*^ mutants cannot be due to poor replicability of the synaptic transmission phenotype. Regardless, both sets of data are consistent with increased spontaneous transmission and a reduction in evoked release.

**Figure 4.**
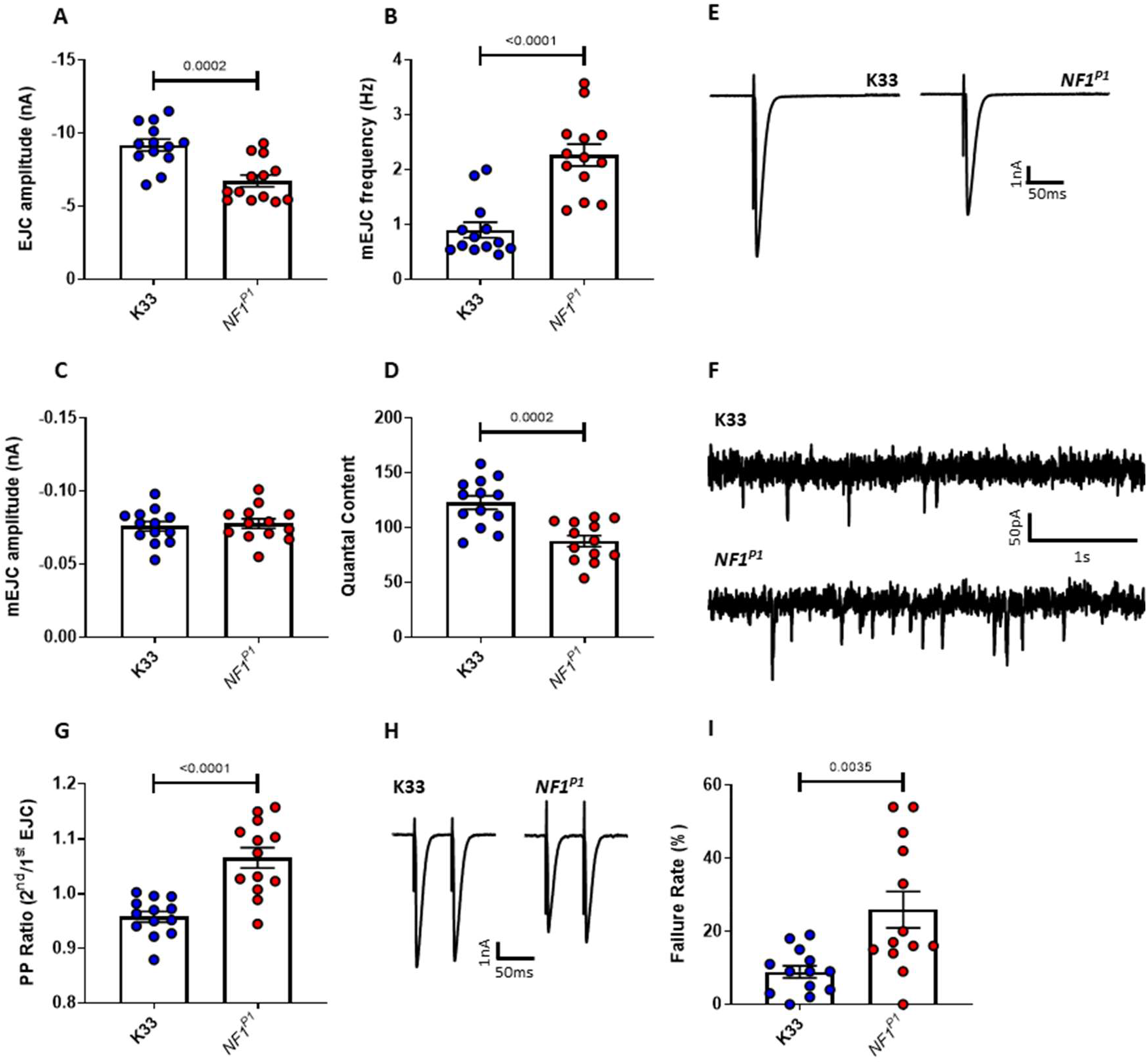
Synaptic current is reduced in *NF1*^*P1*^ mutants. *A)* Under voltage clamp, EJC amplitude is significantly reduced in *NF1*^*P1*^ mutants. *B)* mEJC frequency is significantly increased in *NFI*^*P1*^ larvae, while *C)* there is no significant difference in mEJC amplitude (p=0.6298). *D)* Quantal content is significantly reduced for *NF1*^*P1*^ larvae. *E-F)* Representative traces of EJCs and mEJCs, respectively. *G)* Under voltage clamp in HL3 saline (1.5 mM Ca^2+^), the paired-pulse ratio (PPR; 2^nd^ EJC amplitude/1^st^ EJC amplitude) is significantly increased in *NF1*^*P1*^ larvae. *H)* Representative traces of two EJCs evoked with a 50 ms interval. *I)* In HL3 saline with a reduced Ca^2+^ concentration (0.4 mM Ca^2+^), the rate at which a stimulus fails to evoke an EJP under current-clamp is significantly greater in *NF1*^*P1*^ larvae. All data are presented as mean ± SEM. All statistical comparisons were made via an unpaired, two-tailed student’s *t*-test.

To provide further evidence for the latter, we carried out paired-pulse recordings in voltage clamp mode. If probability of transmitter release is reduced, fewer vesicles will fuse with the presynaptic membrane to release transmitter upon the first stimulus. Therefore, a comparatively higher number of vesicles will be available to fuse with the membrane upon the second stimulus. As a result, the paired-pulse ratio (PPR; see Methods) will be greater.

In *NF1*^*P1*^ larvae, the PPR is indeed significantly increased compared to that of K33 controls (Figure 4G-H). Furthermore, PPR values in K33 controls indicate a prevalence of paired-pulse depression rather than facilitation, as the amplitude of the second EJC is lower than that of the first (demonstrated by a PPR of less than 1). This is to be expected in wildtype larvae at physiological calcium levels, due to the large number of vesicles releasing transmitter upon the initial stimulus. In contrast, paired-pulse facilitation (an increase in EJC amplitude upon the second stimulation) was more common amongst *NF1*^*P1*^ larvae.

We also examined release probability by recording EJPs in the presence of reduced (0.4 mM) Ca^2+^ and calculating the rate at which a stimulus fails to evoke a response (Figure 4I). A significantly greater percentage of stimuli failed to evoke an EJP at the NMJ of *NF1*^*P1*^ larvae compared to that of K33 controls, which is consistent with a reduced probability of transmitter release, and, consequently, a smaller synaptic current.

### Increased post-synaptic R_in_ compensates for reduced evoked release

A reduction in EJC amplitude, despite no change in EJP amplitude, is indicative of increased muscle R_in_ in *NF1*^*P1*^ larvae. This would enable a smaller synaptic current to generate a larger voltage response. For example, mEJP amplitude (recorded under current clamp) is greater in muscle 7 than muscle 6 of wild-type 3^rd^ instar *Drosophila* larvae, despite indistinguishable mEJC amplitudes, owing to the greater R_in_ of the former (Powers et al., 2016). In keeping with this possibility, we observed significantly greater R_in_ values for muscle 6 in *NF1*^*P1*^ larvae compared to K33 controls (Figure 5A-B). Moreover, this increase was also seen following presynaptic (*elav>NF1*^*RNAi*^*;Dicer2*), but not postsynaptic (*MHC>NF1*^*RNAi*^*;Dicer2*) knockdown of *NF1* expression (Figure 5C), consistent with it being a homeostatic response to altered presynaptic signalling, and not a direct result of cell-autonomous loss of *NF1*.

**Figure 5.**
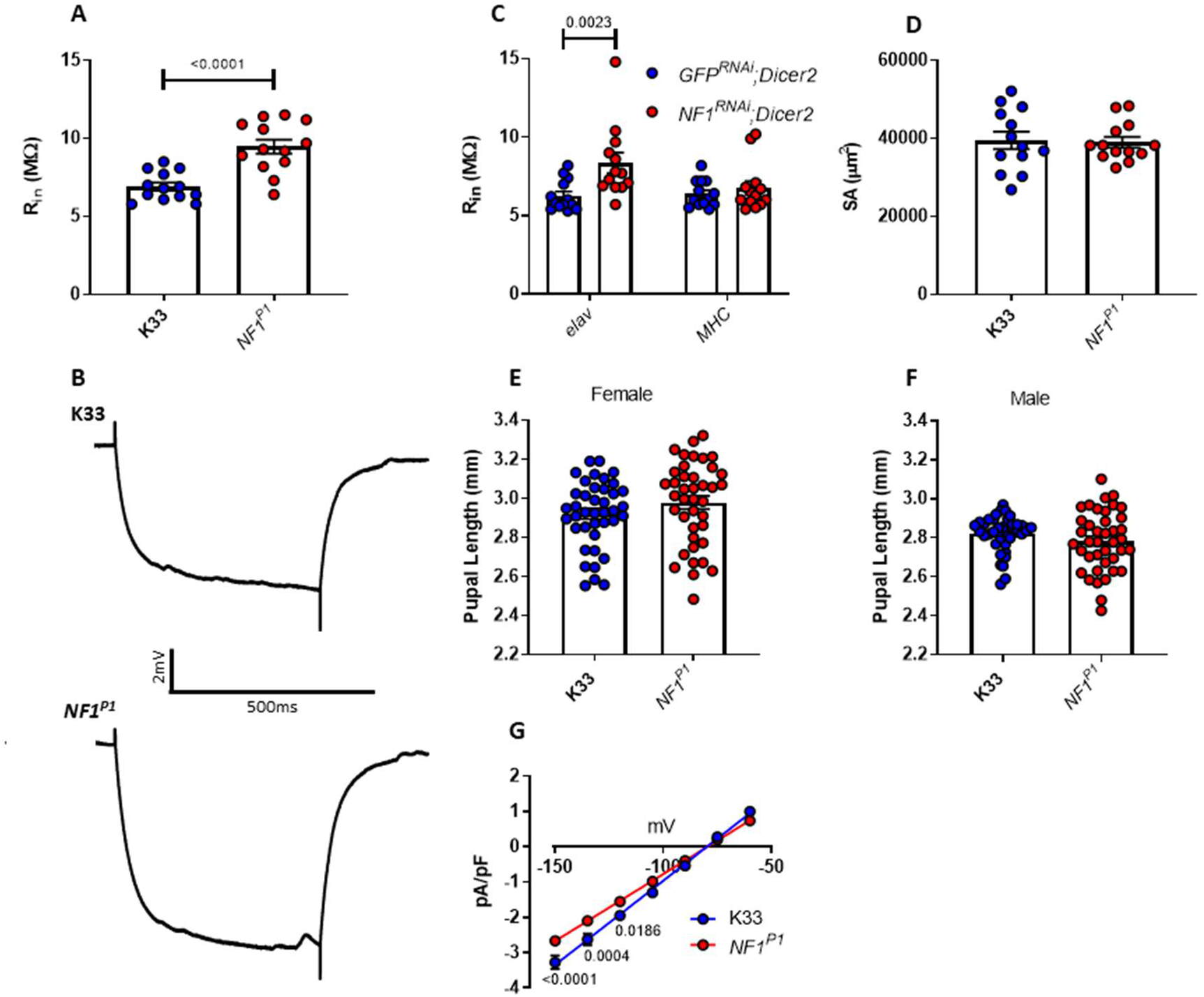
An increase in postsynaptic Rin compensates for reduced evoked transmission in *NF1*^*P1*^ larvae. *A)* Postsynaptic Rin is significantly increased at the *NFI*^*P1*^ NMJ. *B)* Representative traces of the voltage response to injection of -1 nA current into the muscle, which was used to estimate the amplitude of Rin, for each genotype. *C)* A significant increase in postsynaptic Rin is seen at the NMJ of larvae following presynaptic (*elav*), but not postsynaptic (*MHC*; p=0.78), knockdown of *NF1*, consistent with it being a homeostatic response to reduced synaptic drive. *D)* Muscle surface area is not significantly different between genotypes (p=0.85). *E-F)* There is no significant difference in pupal length between females (p=0.18) or males (p=0.19), respectively, of the two genotypes. *G)* IV plot of leak currents as measured under voltage clamp from a holding potential of -80 mV. Current has been normalised to capacitance (pA/pF) to account for possible differences in muscle size. The slope of the *NF1*^*P1*^ linear regression is significantly different (p<0.0001) to that of K33, as are pA/pF values at – 150 mV, -135 mV and –120 mV. All data are presented as mean ± SEM. All data sets were statistically compared via an unpaired, two-tailed student’s *t*-test except for panels C and G, in which data were analysed via two-way ANOVA followed by Sidak’s post-hoc test, to compare *C) NF1*^*RNAi*^*;Dicer2* and *GFP*^*RNAi*^*;Dicer2* larvae within each driver group or *G) NF1*^*P1*^and K33 larvae at each voltage step.

An increase in R_in_ may come about by either a decrease in muscle size or a reduction in leak current, a voltage-independent current that is present at rest and important in regulating membrane excitability. While reduced muscle size would seem plausible, given that a pupal growth defect is one of the most frequently observed phenotypes in *Drosophila* lacking *NF1* expression (The et al., 1997; Walker et al., 2013; Walker et al., 2006), we observed no significant change to muscle surface area (segment A3) in *NF1*^*P1*^ larvae (Figure 5D). Moreover, we were unable to discern any significant difference in pupal length between *NF1*^*P1*^ and K33 larvae for either females (Figure 5E) or males (Figure 5F). However, there was a significant reduction in the amplitude of the leak current in muscle 6 of *NF1*^*P1*^ larvae at more hyperpolarised potentials, as well as a significant difference in the slope of the linear regression (Figure 3G), which would imply a reduction in leak current at more depolarised potentials as well.

### Reduced *NF1* expression in cholinergic neurons is sufficient to induce neuronal hyperexcitability and tactile hypersensitivity

Our consistent observations of an increase in spontaneous release is indicative of an enhancement of neuronal excitability. Should loss of *NF1* induce hyperexcitability within the CNS, as well as at the NMJ, this could incur downstream consequences on larval behaviour. Therefore, we passively recorded endogenous activity from muscle 6 in semi-intact larval preparations in which the CNS had not been removed.

Typically, neuronal activity occurred in firing ‘bursts’ (defined in Methods). We observed a significant increase in time spent burst firing in *NF1*^*P1*^ larvae over a 5 min period (Figure 6A-D). This was due to a significant increase in the number of individual bursts, rather than a change in mean burst duration. Furthermore, some traces (*n* = 3) for *NF1*^*P1*^ larvae displayed almost continuous firing across the recording period (lower *NF1*^*P1*^ trace, Figure 6D), with individual bursts not obviously distinguishable from each other. In contrast, several K33 traces (*n* = 4) showed a complete absence of activity (lower K33 trace, Figure 6D). Thus, it appears that loss of *NF1* does indeed lead to hyperexcitability of moto-neurons of *Drosophila* larvae.

**Figure 6.**
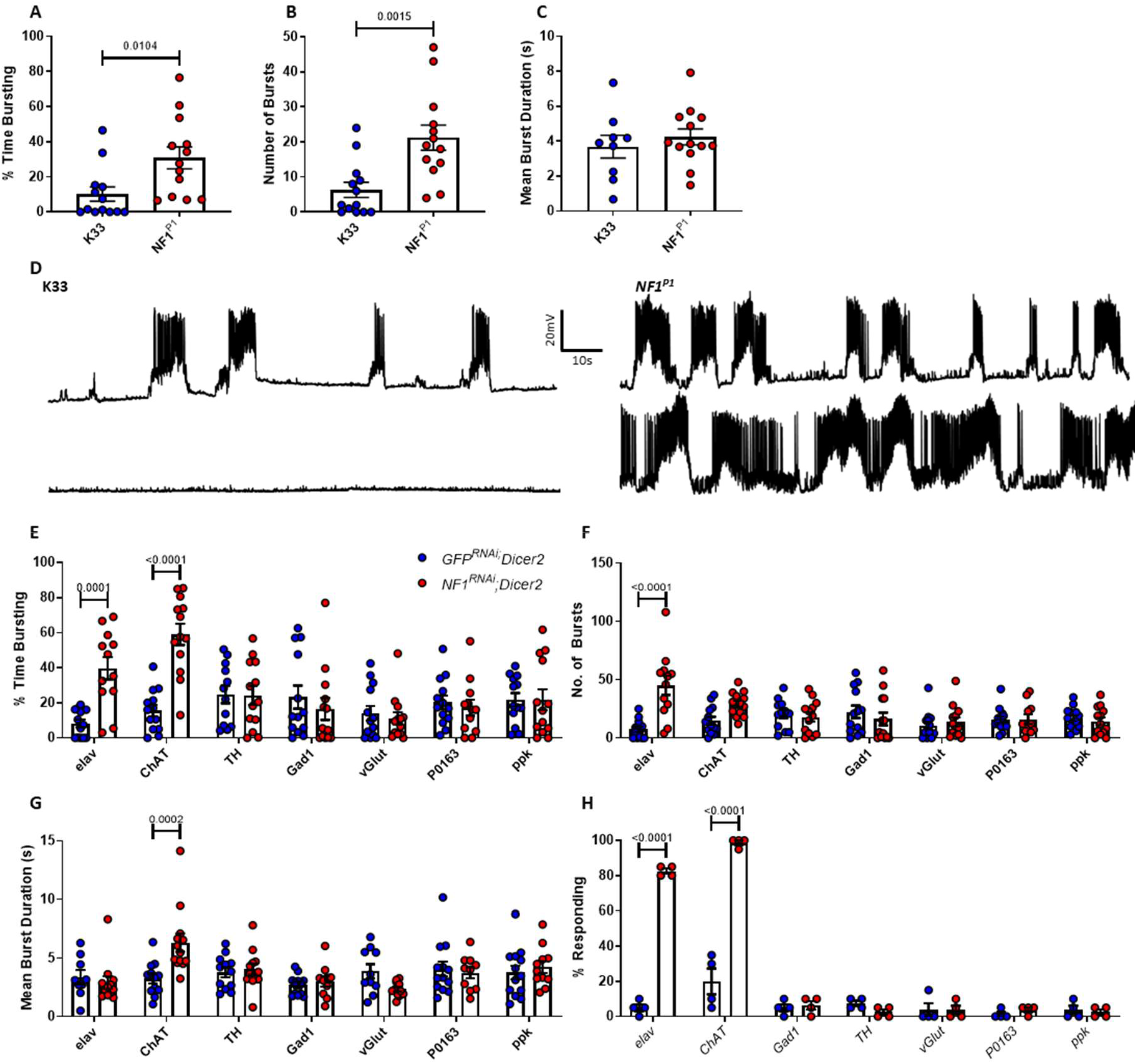
Loss of *NF1* in cholinergic neurons results in neuronal hyperexcitability and tactile hypersensitivity. *A–C)* The percent time spent burst firing over a 5-minute period was significantly greater for *NF1*^*P1*^ mutants compared to K33 controls, as was the number of individual bursts. In contrast, mean burst duration was unchanged between genotypes (p=0.48). *D)* Representative traces of burst firing for data in A-C. *E) elav-* driven knockdown of *NF1*^*RNAi*^*;Dicer2* induces a significant increase in percent time bursting, as does *ChAT-GAL4*. In contrast, *TH-* (p>0.99), *Gad1-* (p=0.94), *vGlut-* (p=0.99), *P0136-* (p=0.99) and *ppk-*driven (p>0.99) knockdowns do not. *F)* As observed in *NF1*^*P1*^ larvae, excessive firing in *elav>NF1*^*RNAi*^*;Dicer2* larvae arises from an increased number of bursts. While there is an increase in burst number for *ChAT>NF1*^*RNAi*^*;Dicer2* larvae, this is non-significant (p=0.11). *G) ChAT>NF1*^*RNAi*^*;Dicer2* larvae exhibit excessive activity via an augmented mean burst duration, which is not seen in any other line. *H)* Only *elav-* and *ChAT-*driven knockdowns of *NF1*^*RNAi*^*;Dicer2* result in tactile hypersensitivity. All data are presented as mean ± SEM. All statistical analyses in A-C were carried out via unpaired student’s *t*-test. Panels E-H were analysed via two-way ANOVA followed by Sidak’s post-hoc test to compare *GFP*^*RNAi*^*;Dicer2* and *NF1*^*RNAi*^*;Dicer2* lines within each *GAL4* driver group.

To determine whether loss of *NF1* in the CNS is responsible for the excessive endogenous firing we observed, we carried out a series of *NF1* knockdowns targeted to different neuronal subtypes. In accord with the data above, pan-neuronal (*elav*) knockdown of *NF1* led to excessive burst firing due to an increase in burst number, but not burst duration (Figure 6E-G). In subsequent knockdown experiments, only cholinergic (*ChAT*) knockdown of *NF1* caused a significant increase in overall burst firing (Figure 6E), consistent with the deficit arising in excitatory interneurons of the CNS as ∼80% of CNS neurons in *Drosophila* are cholinergic (Lee and O’Dowd, 1999). That knockdown of *NF1* under the control of *GAL4* drivers (*ppk* and *P0163*) that express in peripheral sensory neurons was unable to induce excessive firing further confirms that the deficit is central in origin. Moreover, dopaminergic (*TH*), GABAergic (*Gad1*) or glutamatergic (*vGlut*) knockdown failed to affect neuronal activity (Figure 6E). We also found that the manner in which activity is altered in *ChAT>NF1*^*RNAi*^*;Dicer2* larvae is different to that in *NF1*^*P1*^ and *elav>NF1*^*RNAi*^*;Dicer2* larvae; following cholinergic knockdown of *NF1*, mean burst duration is significantly increased, but the increase in burst number is not statistically significant (Figure 6F-G). Therefore, while loss of *NF1* in excitatory cholinergic neurons of the CNS is sufficient to cause neuronal hyperexcitability, it remains possible that the exact nature of this phenotype may be shaped by loss of *NF1* in other cell populations.

Next, we investigated whether neuronal hyperexcitability correlates with hypersensitivity to mechanical stimuli. Indeed, only *elav*-and *ChAT*-driven knockdown of *NF1* recapitulated the tactile hypersensitivity phenotype (Figure 6H). While no conclusions on the causal nature of this relationship can be drawn from this data alone, given that, as mentioned above, the large majority of CNS neurons are cholinergic, it is still consistent with the possibility that neuronal hyperexcitability in the CNS underlies hypersensitivity to mechanical stimulation in *NF1*^*-/-*^ mutant larvae.

### Increased Ras signalling underlies impaired synaptic transmission and tactile hypersensitivity

In both vertebrates and invertebrates, *NF1* has been implicated in myriad molecular pathways. The most prominent molecular roles are as a Ras-GAP that functions to inhibit Ras activity by catalysing the hydrolysis of active Ras-GTP to inactive Ras-GDP, and as a positive regulator of cAMP/PKA cascades via the stimulation of adenylyl cyclase. Consequently, loss of *NF1* leads to excessive Ras and diminished cAMP/PKA signalling (Bergoug et al., 2020). Moreover, levels of phosphorylated MAPK have been shown to be augmented in the *NF1*^*P1*^ mutant, indicative of excessive Ras activity (Botero et al., 2021; Williams et al., 2001). Therefore, we investigated which of these biochemical functions, if either, might explain the role of *NF1* in regulating synaptic transmission and larval mechanoreception. We hereafter use spontaneous miniature transmission (i.e. mEJP frequency) as a measure of neuronal excitability (Mosca et al., 2005).

*Drosophila* possess two Ras proteins against which *NF1* has been shown to exert GAP activity: Ras85D (also termed Ras1), which is homologous to human H-Ras, K-Ras, and N-Ras, and Ras64B (also termed Ras2), which is homologous to human R-Ras proteins (Walker et al., 2006). We used pan-neuronal (via *elav-Gal4*) RNAi to attenuate expression of these two proteins in the *NF1*^*P1*^ mutant background. Knockdown of either *Ras64B* (Figure 7A-F) or *Ras85D* (Figure S2A-F) restored mEJP frequency, mEJP amplitude, and quantal content to levels not significantly different from heterozygote controls. Knockdown of either protein also fully rescued the tactile hypersensitivity phenotype (Figure 7G, Figure S2G). That reduced expression of each Ras homolog, alone, is able to rescue synaptic dysfunction and abnormal behaviour may suggest some level of redundancy in the function of each protein. Alternatively, it could indicate that increased activity of both *Ras64B* and *Ras85D* following loss of *NF1* is necessary to alter synaptic transmission and behaviour, such that reduced expression of either homolog is sufficient to restore these to normal.

**Figure 7.**
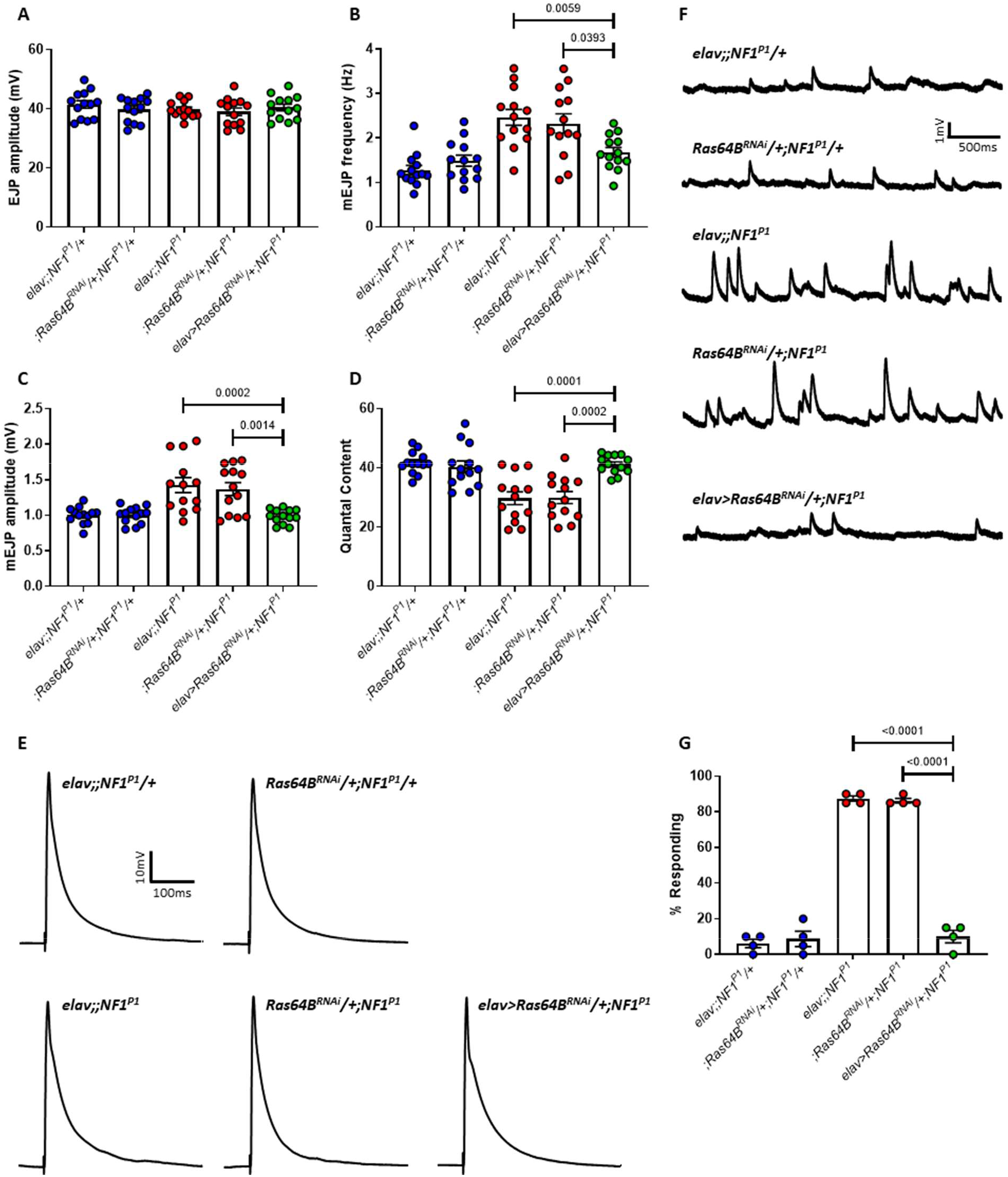
Knockdown of *Ras64B* rescues synaptic transmission deficits and tactile hypersensitivity in *NF1*^*P1*^ larvae. *A)* EJP amplitude was not significantly different between any of the lines tested. *B)* mEJP frequency in *elav>Ras64B*^*RNAi*^*/+;NF1*^*P1*^ larvae (rescue line; green circles) is significantly reduced compared to both homozygous mutant lines (red circles). Expression of UAS-*Ras64B*^*RNAi*^ also rescues *C)* increased mEJP amplitude and *(D)* reduced quantal content. There were no significant differences between the rescue line and either heterozygous control (blue circles) for any parameter examined. Furthermore, in panels B-D, both heterozygous controls were significantly different to both homozygous mutant controls, and there were no significant differences between either of the heterozygous controls or either of the homozygous mutant controls, respectively. *E–F)* Representative traces of EJPs and mEJPs, respectively, for each of the lines tested in A-D. *G)* Pan-neuronal expression of *UAS-Ras64B*^*RNAi*^ is sufficient to rescue tactile hypersensitivity in *NF1*^*P1*^ larvae. All data are presented as mean ± SEM. All statistical comparisons were made via one-way ANOVA followed by Tukey’s multiple comparisons test.

Next, we examined whether diminished cAMP activity might contribute to the signalling deficit. Driving expression of a constitutively-active UAS-PKA* under the control of *elav-GAL4* is lethal (Walker et al., 2013), and so was not a viable option for upregulating cAMP/PKA signalling here. Therefore, we raised larvae on fly food supplemented with the cell-permeable cAMP analogue db-cAMP in an attempt to pharmacologically raise cAMP levels. The concentration administered (10 µM) was previously shown to rescue structural abnormalities at the *NF1*^*E2*^ mutant larval NMJ (Tsai et al., 2012). However, this treatment did not rescue mEJP frequency, mEJP amplitude, or quantal content in *NF1*^*P1*^ mutants (Figure S3A-F), nor did it affect larval behaviour (Figure S3G). Together, our data strongly implicate Ras signalling pathways in *NF1*-mediated regulation of synaptic transmission, and suggest that this occurs independently of the role of *NF1* in cAMP/PKA activation.

## Discussion

Approximately 60% of children with ASD display clear differences in tactile sensitivity, which may, for example, manifest as discomfort when grooming, a negative reaction to being touched, or a dislike of standing close to others (Tomchek and Dunn, 2007). Adults with ASD also exhibit hypersensitivity to stimuli to a degree that is correlated with their Autism Quotient score (Tavassoli et al., 2014). Indeed, sensory processing impairments in early development are predictive of later ASD diagnosis and may even directly contribute to other ASD traits (Robertson and Baron-Cohen, 2017). Thus, models of behavioural hypersensitivity are appropriate for the investigation of core ASD symptom aetiology.

Here, we demonstrate tactile hypersensitivity in two different *NF1*^*-/-*^ larval lines that is specific to loss of *NF1* function. While the method of stimulation used here differs from that in other studies of larval nociception (Mauthner et al., 2014; Walcott et al., 2018), we deem it appropriate for our purposes given that it generally does not induce a response in control larvae and is thus suitable for identifying mutations that give rise to tactile hypersensitivity. Moreover, due to the strength and consistency of the phenotype, as well as the ease with which it can be characterised, we expect that it will provide a valuable output for the screening of genetic and pharmacological modifiers of *NF1*. It also strengthens the face validity of *Drosophila* NF1 models and lends additional support for their application to ASD research. Although tactile sensitivity, in particular, has not been investigated in individuals with NF1 and ASD, a recent study has shown that auditory processing abnormalities present in early development correlate with the later emergence of ASD traits in NF1 infants (Begum-Ali et al., 2021).

Consistent with a belief that synaptic transmission abnormalities contribute to ASD pathophysiology, loss of *NF1* expression under physiologically relevant conditions (Stewart et al., 1994) results in a reduction in evoked release that is homeostatically compensated for, but an increase in spontaneous release. Despite observing similar phenotypes across four different NF1 paradigms here (Figures 2 and 3, Figure S1), our findings are, nevertheless, inconsistent with those from a previous study (Tsai et al., 2012) in which it was shown that *NF1*^*E2*^ mutants display an increased EJP amplitude and quantal content, with no change in mEJP amplitude or frequency. The reason for this discrepancy is unclear. One possible explanation was that the study by Tsai et al. (2012) used reduced Ca^2+^ (0.2 mM) and Mg^2+^ (4 mM). However, we were unable to replicate their findings even under identical conditions (data not shown), and instead observed a milder version of the phenotypes shown in Figure 2. That we see a similar phenotype in different *NF1* mutant and knockdown lines, each with a different genetic background, also precludes mutation-dependent mechanisms and/or genetic modifiers as likely explanations. Therefore, in the absence of any clear effector, we postulate that some environmental factor may be involved, such as the diet on which the larvae were raised. In support of this, it was recently shown that two different fly food recipes, both of which are considered standard, may enhance or suppress the severity of a seizure phenotype of mutant *para*^*Shu*^ flies (Kasuya et al., 2019).

There are also notable differences between the findings presented here and those from murine models of NF1. In *NF1* mutant hippocampal neurons, evoked release, whether inhibitory (Omrani et al., 2015) or excitatory (Wang et al., 2010), is augmented, in contrast to the data presented here showing an overall reduction in evoked release. Furthermore, while the frequency of spontaneous and/or miniature transmission is typically found to be increased in NF1 mouse models, in the hippocampus (Cui et al., 2008; Omrani et al., 2015), medial prefrontal cortex, and striatum (Shilyansky et al., 2010), this is the case only for inhibitory and not excitatory currents. Although an increase in mEPSC frequency is present in the basolateral amygdala of *NF1*^*+/-*^ mice, the molecular mechanism giving rise to this is unclear, as the deletion of *PAK1*, which would be expected to attenuate Ras/MAPK signalling, leads to a further increase in mEPSC frequency (Molosh et al., 2014). Conversely, our data suggest that synaptic dysfunction at the *NF1*^*P1*^ NMJ is a direct result of excessive Ras activity, in line with studies of increased inhibitory transmission in NF1 (Cui et al., 2008; Molosh et al., 2014; Omrani et al., 2015; Shilyansky et al., 2010).

Studies of NF1 mouse models also do not find an increase in the amplitude of mIPSCs or mEPSCs (Cui et al., 2008; Molosh et al., 2014; Omrani et al., 2015). Similarly, mEJC amplitude in *NF1*^*P1*^ larvae is unaltered, and we posit that the increase in mEJP amplitude is not a result of the primary deficit but rather reflects a compensatory response to reduced evoked release such that EJP amplitude is unaltered. This homeostatic response appears to be a non-cell-autonomous postsynaptic increase in R_in_, mediated by a reduction in muscle leak current, and not due to any change in muscle size (Figure 5). That EJP amplitude is marginally reduced in *elav>NF1*^*RNAi*^*;Dicer2* larvae may indicate that this response does not occur to the same extent in this line as in *NF1*^*P1*^ or *NF1*^*E2*^ larvae.

Differences between our data and those from murine models are not necessarily unexpected, given that discrepancies also exist between mouse and human clinical studies. For example, *NF1*^*+/-*^ mice display an increase in GABA/Glu ratio in the prefrontal cortex and striatum, and a significant increase in GABA_A_ receptor expression in the hippocampus (Goncalves et al., 2017), all consistent with electrophysiological studies indicating enhanced inhibitory firing (Cui et al., 2008; Molosh et al., 2014; Omrani et al., 2015; Shilyansky et al., 2010). In contrast, in individuals with NF1, GABA levels were found to be significantly reduced in the prefrontal and visual cortex, with a reduction in GABA_A_ receptor density in the striatum (Ribeiro et al., 2015; Violante et al., 2016; Violante et al., 2013). These differences may suggest that *NF1* has numerous molecular and cellular functions that may be specific to a particular neuronal subtype or brain region, and which may account for apparent discrepancies between studies of *NF1* in diverse model systems. It is also possible that these differences reflect homeostatic alterations to early changes in neuronal activity, rather than the primary deficit (Nelson and Valakh, 2015).

An important question remaining to be resolved is how an increase in spontaneous transmission can be reconciled with a reduction in evoked transmission. One possibility is that *NF1* is involved in the ‘clamp’ that prevents vesicle fusion in the absence of stimulation, thereby restricting spontaneous release. This would result in spontaneous release being increased at the expense of evoked transmission, as is seen in *synaptotagmin* mutant larvae (Littleton et al., 1994; Littleton et al., 1993). However, in these mutants, the degree of dysfunction is more severe and is not compensated for, such that evoked EJP amplitude is significantly reduced. Alternatively, because spontaneous and evoked release at the NMJ do not necessarily occur at the same active zones (AZs), with certain AZs only displaying one form of release (Melom et al., 2013), *NF1* may play a role in determining the type of release in which a particular AZ is involved. If *NF1* promotes formation and/or function of ‘evoked release’ AZs, its loss of expression might be predicted to lead to fewer evoked release events and an increase in spontaneous events. Other possibilities include enhanced spontaneous release, rather than being a direct result of reduced *NF1* expression, being a consequence of the homeostatic increase in muscle R_in_ and subsequent increase in neuronal excitability, assuming a similar mechanism occurs in synapses between neurons as well as at the NMJ. Clearly future studies will be required to distinguish between these possible mechanisms.

The increase in mEJP frequency we observe is indeed consistent with neuronal hyperexcitability. This lends support to the excitatory/inhibitory imbalance theory of ASD, which posits that net excitatory activity is increased in certain neural systems such as those involved in social and sensory processing (Rubenstein and Merzenich, 2003). The deficit we observe includes central neurons, as only cholinergic knockdown of *NF1* recapitulated this phenotype, whilst knockdown in peripheral/sensory neurons did not. Likewise, only cholinergic knockdown of *NF1* gave rise to tactile hypersensitivity, whereas knockdown in sensory/peripheral neurons did not (Figure 6F). This is important in supporting the relevance of our work to ASD. Initially, an argument could have been made that the enhanced nocifensive response is more appropriate as a model of severe pain, another common complication in NF1. However, research into pain in NF1 strongly points to a role for neuronal hyperexcitability in peripheral sensory neurons (Bellampalli and Khanna, 2019), suggesting that this behaviour in *NF1*^*-/-*^ larvae is unrelated.

Cortical hyperexcitability in ASD is often suggested to underlie inappropriate neural responses to stimuli and behavioural hypersensitivity (Takarae and Sweeney, 2017). Notably, tactile stimulation leads to a hyperexcitable response in the primary somatosensory cortex of a mouse model of the ASD-associated Fragile X Syndrome and is associated with pyramidal dendrite hyperexcitability (Zhang et al., 2014). Therefore, we speculate that excessive excitatory firing, arising from the central locomotor circuitry, underlies tactile hypersensitivity in *NF1*^*P1*^ larvae. In support of this, both increased spontaneous transmission, which is typically indicative of neuronal hyperexcitability (Mosca et al., 2005), and tactile hypersensitivity are rescuable via the knockdown of Ras proteins, consistent with the two phenotypes sharing a common mechanism. However, our data are only correlational, and narrowing down the neuronal subtypes involved, as well as the downstream Ras effectors, will be necessary to validate this hypothesis.

Finally, it is interesting to consider how synaptic dysfunction and tactile hypersensitivity might relate to other ASD-relevant phenotypes in *Drosophila* models of NF1. Recently, it was shown that *NF1*^*P1*^ male flies exhibit social interaction deficits in the form of impaired courtship behaviour (Moscato et al., 2020). However, this requires *NF1* in peripheral *ppk23*-expressing chemosensory neurons (Moscato et al., 2020), thus is unlikely to be related to larval hypersensitivity. On the other hand, excessive grooming behaviour in *NF1*^*P1*^ flies, assumed to reflect the repetitive behaviours characteristic of ASD individuals (King et al., 2016), arises from loss of *NF1* in cholinergic neurons of the CNS and is Ras-dependent (King et al., 2020). Although excessive grooming arises due to developmental loss of *NF1* specifically in the pupal stage (King et al. 2020), it is still possible that the underlying mechanisms for both are similar. Moreover, degree of hypersensitivity has been shown to positively correlate with repetitive behaviour severity in ASD children (Boyd et al., 2010).

We have shown in this study that *NF1*^*-/-*^ larvae display hypersensitivity to a mechanical stimulus and exhibit excitatory synaptic transmission deficits suggestive of neuronal hyperexcitability, both of which strengthen the argument for the use of *Drosophila* models of NF1 in the study of ASD. Although the data presented here are consistent with a causal relationship between the two phenotypes, further evidence will be required to conclusively demonstrate this. We predict that these phenotypes will provide tractable outputs by which to screen for genetic and/or therapeutic modifiers of NF1 and/or ASD in the future.

## Supporting information

Video S1. Typical K33 response to mechanical stimulation

Video S2. Typical NF1P1 response to mechanical stimulation

## Acknowledgements

This work was funded by a Medical Research Council Doctoral Training Partnership to A.D., and by funding from the Biotechnology and Biological Sciences Research Council (BB/L027690/1) to R.A.B. We thank Seth Tomchik for providing *NF1*^*P1*^ and K33 lines, Cheng-Ting Chien for providing *NF1*^*E2*^ and *w*^*1118*^ lines, and James Walker for providing the *UAS-NF1* line. Work on this project benefited from the Manchester Fly Facility, established through funds from the University and the Wellcome Trust (087742/Z/08/Z). S.G. is supported by the Neurofibromatosis Therapeutic Acceleration Program (NTAP) Francis Collins Scholarship. D.G.E. is supported by the Manchester NIHR Biomedical Research Centre (IS-BRC-1215-20007).

## Author Contributions

R.A.B., S.G. and D.G.E. conceptualised and supervised the project. A.D. carried out all experimental work and data analysis, which was overseen by R.A.B. The manuscript was written primarily by A.D., with comments and corrections from R.A.B. and S.G.

## Methods

### Fly lines and maintenance

*NF1*^*P1*^ and K33 fly lines were obtained from Dr. Seth Tomchik (Scripps Research Institute, Florida, USA). The *NF1*^*P1*^ mutation was generated by mobilisation of a P-element, resulting in total deletion of the *NF1* gene except for exon 1 and complete ablation of *NF1* expression (The et al., 1997). The K33 line is the parental line containing the P-element used to generate the deletion (The et al., 1997), and is frequently used as a control against *NF1*^*P1*^ (Buchanan and Davis, 2010; Guo et al., 2000; Moscato et al., 2020). Both lines have been back-crossed into the *w*^*CS10*^ background such that K33 functions as an isogenic control. The *NF1*^*E2*^ line, which contains an EMS-induced nonsense mutation within the *NF1* gene (Walker et al., 2006), and its isogenic *w*^*1118*^ control line (Tsai et al., 2012) were obtained from Dr. Cheng-Ting Chien (Institute of Molecular Biology, Academia Sinica, Taiwan). The *UAS-NF1* line used here (King et al., 2020) was obtained from Dr. James Walker (Centre for Genomic Medicine, Harvard Medical School, USA). For *NF1* knockdowns via RNA interference (RNAi), *UAS-NF1*^*RNAi*^ transgenes were combined with *UAS-Dicer2* to augment knockdown efficacy, and *GAL4* driver lines were crossed to either *UAS-NF1*^*RNAi*^;*UAS-Dicer2* (experiment) or *UAS-GFP*^*RNAi*^*;UAS-Dicer2* (control). *GAL4* driver lines and their sources are as follows: *elav*^*c155*^*-GAL4* (Bloomington, #458), *MHC-GAL4* (Baines Lab), *ChAT:BAC-GAL4* (gift from Dr. Steven Stowers), *TH-GAL4* (Bloomington, #8848), *Gad1-GAL4* (gift from Dr Matthias Landgraf), *vGlut*^*OK371*^*-GAL4* (Bloomington, #26160), *P0163-GAL4* (Kyoto, #103168), *ppk-GAL4* (Bloomington, #32079). VDRC *UAS-NF1*^*RNAi*^ line #109637 was used for all *NF1* knockdown experiments except those in Figure S1, for which construct #35877 was used. For rescue experiments involving the pan-neuronal expression of *UAS-NF1, UAS-Ras64B*^*RNAi*^ (VDRC #110574), or *UAS-Ras85D*^*RNAi*^ (VDRC #106642), these transgenes were combined into the *NF1*^*P1*^ mutant background and crossed to *elav*^*c155*^*-GAL4*;;*NF1*^*P1*^. Control lines for rescue experiments comprised each parental line crossed to either *NF1*^*P1*^ (producing homozygous mutant controls) or K33 (producing heterozygous controls that do not display any phenotype relative to K33; data not shown). All lines and crosses were maintained on standard cornmeal medium in a 12:12 light:dark cycle at 25°C.

### Larval mechanoreception

Water (2 ml) was added to a 35 mm petri dish containing Silgard into which wall-climbing third instar larvae were placed. An Austerlitz Minutiens stainless steel insect pin (diameter = 0.1 mm), cut to approximately 2 mm in length and held horizontally between a pair of forceps, was pressed down firmly upon the posterior end of the larva (Figure 1A, Videos S1 and S2). Care was taken to ensure that the pin was applied with the same pressure to each larva, and all repeats of the assay were carried out by the same experimenter, blinded to genotype and/or treatment. Whether or not the larva exhibited a stereotypic rolling motion, characteristic of the nocifensive response within 10 seconds of stimulation, was noted. Only a full 360° roll was classified as a nocifensive response. For each genotype and/or treatment group, 4 trials of 20 larvae were carried out, from which mean percentage responding larvae was calculated. For experiments using progeny derived from transgenic parental lines (e.g. RNAi experiments), each individual trial comprised larvae taken from separate, independent genetic crosses.

### Electrophysiology

#### Saline and recording criteria

To examine synaptic transmission at the larval NMJ, wall-climbing third instar larvae were dissected in HL3 saline without CaCl_2_ (NaCl, 70mM; KCl, 5mM; MgCl_2_ hexahydrate, 20mM; NaHCO_3_, 10mM; sucrose, 115mM; trehalose, 5mM; HEPES, 5mM; pH=7.25, adjusted with NaOH) to expose ventral body wall muscles, before the CNS was removed to permit access to the peripheral nerves (except when recording endogenous excitatory activity; see *Semi-intact recordings*). Larval preparations were then washed 5 times in HL3 saline containing the Ca^2+^ concentration in which the experiment was to be carried out. This was 1.5 mM (Stewart et al., 1994) for all experiments except those characterising EJP failure rate, for which 0.4 mM was used. All recordings were performed at room temperature and taken from muscle 6 in segments A2 - A4. Recording and suction pipettes were pulled from thick-wall borosilicate capillaries with filament (GC100F-10, Harvard Apparatus, UK) using a Flaming/Brown Micropipette Puller, model P-97 (Sutter Instrument, USA). Recording pipettes were pulled to a resistance of 20 - 35 MΩ when filled with 3 M KCl, while current-passing pipettes (for two-electrode voltage clamp; TEVC) were pulled to a resistance of 15 – 25 MΩ when filled with 3 M KCl. Suction pipettes (for nerve stimulation) were broken and heat-polished to the final desired size then filled with HL3 saline + CaCl_2_. Immediately prior to insertion of the recording electrode into the muscle, pipette offset was corrected to 0 mV. Recordings were taken if R_in_, measured as the voltage response to injection of -1 nA hyperpolarizing current, following electrode insertion into the muscle exceeded 5 MΩ (with exception; see *Semi-intact recordings*) and resting membrane potential (V_m_) was at or below -60 mV. Recordings were also only accepted for data analysis if the voltage drift after the recording electrode was removed from muscle did not exceed ±5 mV. Voltage responses were recorded in current-clamp mode using an Axopatch 200B microelectrode amplifier, Digidata 1322A, and Clampex 10.3 data acquisition software (Molecular Devices, CA, USA). All recordings and subsequent analyses were carried out by an experimenter blinded to genotype and/or drug treatment. Except for recordings in which the CNS was left *in situ* (see *Semi-intact recordings*), *n* = 13 for all electrophysiology experiments.

#### Current-clamp recording of (m)EJPs

EJPs were evoked via 0.2 ms stimulation of the nerve using a DS2A Isolated Voltage Stimulator (Digitimer Ltd., UK). Each stimulation used the minimum voltage necessary to stimulate both Ib and Is motor neuron inputs to generate EJPs of consistent amplitude. Ten EJPs were recorded per muscle at a stimulation frequency of 0.5 Hz. From the same muscle, the membrane potential (V_m_) of the muscle was then recorded for two minutes in the absence of stimulation to obtain mEJPs. No more than 2 sets of recordings (evoked EJP recording followed by passive mEJP recording) satisfying all acceptance criteria listed above were taken from any one larval preparation. Mean EJP amplitude and resting V_m_ were calculated on Clampfit 10.3 analysis software (Molecular Devices). Mean mEJP amplitude and mEJP frequency were calculated using Mini Analysis (Synaptosoft Inc. GA, USA). All data were then exported to Microsoft Excel, and amplitudes corrected for differences in resting V_m_ by applying the equation:

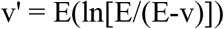

where v’ is the corrected EJP amplitude, v is the recorded EJP amplitude, and E is the driving force which, assuming a reversal potential of 0mV, is equal to resting V_m_ (Feeney et al., 1998). Quantal content was calculated by dividing the corrected mean EJP amplitude by the corrected mean mEJP amplitude.

#### TEVC recordings of (m)EJCs

In addition to the criteria specified for current-clamp experiments, recordings in TEVC were carried out if, following insertion of the current-passing electrode into the muscle after the recording electrode, resting V_m_ was depolarised no more than –50 mV and voltage readings from each electrode were within 5mV of each other. To record EJCs and mEJCs, V_m_ was held at -70 mV, and a 1 kHz filter was applied to facilitate the identification of mEJCs from baseline during analysis. Clampfit 10.3 analysis software (Molecular Devices) was used to calculated mean EJC amplitude, mean mEJC amplitude, and mEJC frequency. Quantal content was calculated by dividing the mean EJC amplitude by the mean mEJC amplitude. The ‘Template Search’ function was used to detect mEJCs, and any events that appeared to be noise were manually excluded. Baseline noise was reduced with a low-pass boxcar filter.

#### Paired-pulse recordings

Paired-pulse experiments were carried out under TEVC. Two stimuli (0.2 ms, 20 Hz) were applied to the nerve 5 times, with 10 seconds between each sweep. The mean amplitude of each EJC was measured from its peak to the baseline V_m_ prior to the first stimulus. The PPR was calculated by dividing the amplitude of the second EJC by that of the first EJC.

#### Failure rate

To calculate EJP failure rate, 100 stimuli (0.2 ms, 0.2 Hz) were applied in HL3 saline containing 0.4 mM CaCl_2_, such that some stimuli failed to evoke an EJP. A failure was defined when no clear depolarisation of the muscle occurred following stimulation, irrespective of amplitude. To remove the possibility that any variation in failure rate may be due to improper nerve stimulation, all nerves were stimulated with 4 V which, based on the stimulation amplitudes required for initial EJP recordings, was sufficient to activate both Ib and Is inputs.

#### Leak current recording

Leak currents were recorded using a protocol similar to that used previously to characterise the *Drosophila* K^+^ leak channel *ORK1* expressed in *Xenopus* oocytes (Goldstein et al., 1996). V_m_ was held at -80mV before application of a step protocol (range: -150 mV to -60 mV; increment: +15 mV; step duration: 75 ms; inter-pulse interval: 1 s). The step range was selected to avoid membrane depolarisation and subsequent muscle contraction, which could disturb the recording electrode. Following each step, V_m_ was hyperpolarised to -150 mV for 15 ms before being returned to -80 mV. All currents were normalised to capacitance to account for possible differences in muscle size and data then fitted with a simple linear regression (GraphPad Prism).

#### Semi-intact recordings

To examine spontaneous EJP generation, muscle V_m_ was recorded passively for 5 minutes in preparations in which the CNS remained *in situ*. Prior to recording, preparations were bathed in saline containing nifedipine (75 μM, 0.3 % DMSO) for 10 minutes to suppress muscle contractions (Kratschmer et al., 2021). Recordings were taken only if contractions were observed at the start of the 10 minute incubation in order to ensure that the CNS was indeed intact, and therefore that lack of endogenous activity did not reflect damage to the CNS incurred during dissection. The frequency of synaptic activity in these experiments meant that measurement of R_in_ as described was not necessarily feasible; therefore, we did not take this into account when deciding whether to proceed with recording (Kratschmer et al., 2021). We defined a burst as starting when ≥5 EJPs occurred within one second (i.e. mean frequency of 0.2 Hz) and as ending when one second or more passed in which ≥5 EJPs were not present, criteria similar but not identical to those previously used (Kratschmer et al., 2021). Traces were analysed using Clampfit 10.3, in which the time spent burst firing at each NMJ, over a 5-minute period, was calculated. We also calculated the number of bursts per recording, and the mean burst duration (total time burst firing / number of bursts). Traces in which we could not clearly identify when both Ib and Is motor neurons were active were excluded from analysis, such that *n* = 12 – 13 for each genotype.

### Drug Treatment

Dibutyryl-cAMP (db-cAMP; Merck Life Science) was dissolved in H_2_O as a 20 mM stock solution. This was then added into molten fly food (<60 °C) to a final concentration of 10 µM, on which larvae were raised throughout development. An equivalent volume of vehicle was added to fly food for controls. To avoid degradation of the compound, the stock solution was stored at -20 °C between uses and disposed of after 1 month.

### Larval measurements

#### Muscle Fibre measurements

Third instar larvae were fillet-dissected to expose muscle 6. Images were captured using a Leica DM6000B microscope. Length and width of segment A3 were obtained with MetaMorph software (version 7.8.13.0, Molecular Devices) via the ‘Region Measurements’ function, from which surface area was then calculated.

#### Pupal length measurements

40 pupae of each sex were collected per genotype. Males and females were identified by the presence or absence of sex combs, respectively, which are clearly visible beneath the pupal casing. Images were acquired using a Leica MZ10F stereo microscope together with Leica Application Suite software (version 4.0.0, Leica Microsystems). Anterior to posterior measurements were then calculated on ImageJ 1.53e (NIH). As wildtype females are typically larger than males, each sex was analysed separately.

### Statistical Analysis

All statistical analysis was carried out in GraphPad Prism (version 8.4.3). Statistical tests used to analyse each data set are indicated in the accompanying figure legend. Experiments comprising only two data sets were analysed via an unpaired, two-tailed student’s *t*-test, while those with three or more ungrouped sets were analysed using a one-way ANOVA followed by Tukey’s post-hoc test. For experiments involving grouped data sets in which data were only compared within groups (i.e. *GAL4* lines each crossed to two RNAi lines), comparisons were made via a two-way ANOVA followed by Sidak’s post-hoc test. For experiments involving grouped data sets in which data were also compared between groups (i.e. K33 and *NF1*^*P1*^ lines each treated with vehicle and db-cAMP), comparisons were made via a two-way ANOVA followed by Tukey’s post-hoc test. For clarity, only relevant, statistically significant comparisons (p<0.05) are displayed within the figures. P values for comparisons relevant to interpreting the data that were not significantly different are given in the figure legend.

## References

American Psychiatric Association. (2013). Diagnostic and Statistical Manual of Mental Disorders, 5 Edition (American Psychiatric Association).

Baio, J., Wiggins, L., Christensen, D.L., Maenner, M.J., Daniels, J., Warren, Z., Kurzius-Spencer, M., Zahorodny, W., Rosenberg, C.R., White, T., et al. (2018). Prevalence of Autism Spectrum Disorder Among Children Aged 8 Years - Autism and Developmental Disabilities Monitoring Network, 11 Sites, United States, 2014. Mmwr Surveillance Summaries 67, 1–23. 10.15585/mmwr.ss6706a1.

Begum-Ali, J., Kolesnik-Taylor, A., Quiroz, I., Mason, L., Garg, S., Green, J., Johnson, M.H., Jones, E.J.H., Team, S., and Team, E. (2021). Early differences in auditory processing relate to Autism Spectrum Disorder traits in infants with Neurofibromatosis Type I. Journal of Neurodevelopmental Disorders 13, 22. 10.1186/s11689-021-09364-3.

Bellampalli, S.S., and Khanna, R. (2019). Towards a neurobiological understanding of pain in neurofibromatosis type 1: mechanisms and implications for treatment. Pain 160, 1007–1018. 10.1097/j.pain.0000000000001486.

Bergoug, M., Doudeau, M., Godin, F., Mosrin, C., Vallee, B., and Benedetti, H. (2020). Neurofibromin Structure, Functions and Regulation. Cells 9, 2365. 10.3390/cells9112365.

Botero, V., Stanhope, B.A., Brown, E.B., Grenci, E.C., Boto, T., Park, S.J., King, L.B., Murphy, K.R., Colodner, K.J., Walker, J.A., et al. (2021). Neurofibromin regulates metabolic rate via neuronal mechanisms in Drosophila. Nature Communications 12, 4285. 10.1038/s41467-021-24505-x.

Boyd, B.A., Baranek, G.T., Sideris, J., Poe, M.D., Watson, L.R., Patten, E., and Miller, H. (2010). Sensory Features and Repetitive Behaviors in Children with Autism and Developmental Delays. Autism Research 3, 78–87. 10.1002/aur.124.

Brugha, T.S., McManus, S., Bankart, J., Scott, F., Purdon, S., Smith, J., Bebbington, P., Jenkins, R., and Meltzer, H. (2011). Epidemiology of Autism Spectrum Disorders in Adults in the Community in England. Archives of General Psychiatry 68, 459–466. 10.1001/archgenpsychiatry.2011.38.

Buchanan, M.E., and Davis, R.L. (2010). A Distinct Set of Drosophila Brain Neurons Required for Neurofibromatosis Type 1-Dependent Learning and Memory. Journal of Neuroscience 30, 10135–10143. 10.1523/jneurosci.0283-10.2010.

Costa, R.M., Federov, N.B., Kogan, J.H., Murphy, G.G., Stern, J., Ohno, M., Kucherlapati, R., Jacks, T., and Silva, A.J. (2002). Mechanism for the learning deficits in a mouse model of neurofibromatosis type 1. Nature 415, 526–530. 10.1038/nature711.

Cui, Y.J., Costa, R.M., Murphy, G.G., Elgersma, Y., Zhu, Y., Gutmann, D.H., Parada, L.F., Mody, I., and Silva, A.J. (2008). Neurofibromin Regulation of ERK Signaling Modulates GABA Release and Learning. Cell 135, 549–560. 10.1016/j.cell.2008.09.060.

Daston, M.M., Scrable, H., Nordlund, M., Sturbaum, A.K., Nissen, L.M., and Ratner, N. (1992). The protein produce of the neurofibromatosis type-1 gene is expressed at highest abundance in neurons, schwann-cells, and oligodendrocytes Neuron 8, 415–428. 10.1016/0896-6273(92)90270-n.

Feeney, C.J., Karunanithi, S., Pearce, J., Govind, C.K., and Atwood, H.L. (1998). Motor nerve terminals on abdominal muscles in larval flesh flies, Sarcophaga bullata: Comparisons with Drosophila. Journal of Comparative Neurology 402, 197–209.

Fortuna, R.J., Robinson, L., Smith, T.H., Meccarello, J., Bullen, B., Nobis, K., and Davidson, P.W. (2016). Health Conditions and Functional Status in Adults with Autism: A Cross-Sectional Evaluation. Journal of General Internal Medicine 31, 77–84. 10.1007/s11606-015-3509-x.

Garg, S., Green, J., Leadbitter, K., Emsley, R., Lehtonen, A., Evans, D.G., and Huson, S.M. (2013). Neurofibromatosis Type 1 and Autism Spectrum Disorder. Pediatrics 132, E1642–E1648. 10.1542/peds.2013-1868.

Goldstein, S.A.N., Price, L.A., Rosenthal, D.N., and Pausch, M.H. (1996). ORK1, a potassium-selective leak channel with two pore domains cloned from Drosophila melanogaster by expression in Saccharomyces cerevisiae. Proceedings of the National Academy of Sciences of the United States of America 93, 13256–13261. 10.1073/pnas.93.23.13256.

Goncalves, J., Violante, I.R., Sereno, J., Leitao, R.A., Cai, Y., Abrunhosa, A., Silva, A.P., Silva, A.J., and Castelo-Branco, M. (2017). Testing the excitation/inhibition imbalance hypothesis in a mouse model of the autism spectrum disorder: in vivo neurospectroscopy and molecular evidence for regional phenotypes. Molecular Autism 8, 47. 10.1186/s13229-017-0166-4.

Guo, H.F., Tong, J.Y., Hannan, F., Luo, L., and Zhong, Y. (2000). A neurofibromatosis-1-regulated pathway is required for learning in Drosophila. Nature 403, 895–898. 10.1038/35002593.

Gutmann, D.H., Ferner, R.E., Listernick, R.H., Korf, B.R., Wolters, P.L., and Johnson, K.J. (2017). Neurofibromatosis type 1. Nature Reviews Disease Primers 3, 17004. 10.1038/nrdp.2017.4.

Howlin, P., and Moss, P. (2012). Adults With Autism Spectrum Disorders. Canadian Journal of Psychiatry-Revue Canadienne De Psychiatrie 57, 275–283. 10.1177/070674371205700502.

Hwang, R.Y., Zhong, L.X., Xu, Y.F., Johnson, T., Zhang, F., Deisseroth, K., and Tracey, W.D. (2007). Nociceptive neurons protect Drosophila larvae from parasitoid wasps. Current Biology 17, 2105–2116. 10.1016/j.cub.2007.11.029.

Hyman, S.L., Shores, A., and North, K.N. (2005). The nature and frequency of cognitive deficits in children with neurofibromatosis type 1. Neurology 65, 1037–1044. 10.1212/01.wnl.0000179303.72345.ce.

Kasuya, J., Iyengar, A., Chen, H.L., Lansdon, P., Wu, C.F., and Kitamoto, T. (2019). Milk-whey diet substantially suppresses seizure-like phenotypes of para(Shu), a Drosophila voltage-gated sodium channel mutant. Journal of Neurogenetics 33, 164–178. 10.1080/01677063.2019.1597082.

Keshishian, H., Broadie, K., Chiba, A., and Bate, M. (1996). The Drosophila neuromuscular junction: A model system for studying synaptic development and function. Annual Review of Neuroscience 19, 545–575. 10.1146/annurev.ne.19.030196.002553.

King, L.B., Boto, T., Botero, V., Aviles, A.M., Jomsky, B.M., Joseph, C., Walker, J.A., and Tomchik, S.M. (2020). Developmental loss of neurofibromin across distributed neuronal circuits drives excessive grooming inDrosophila. Plos Genetics 16, e1008920. 10.1371/journal.pgen.1008920.

King, L.B., Koch, M., Murphy, K.R., Velazquez, Y., Ja, W.W., and Tomchik, S.M. (2016). Neurofibromin Loss of Function Drives Excessive Grooming in Drosophila. G3-Genes Genomes Genetics 6, 1083–1093. 10.1534/g3.115.026484.

Knapp, M., Romeo, R., and Beecham, J. (2009). Economic cost of autism in the UK. Autism 13, 317–336. 10.1177/1362361309104246.

Kratschmer, P., Lowe, S.A., Buhl, E., Chen, K.F., Kullmann, D.M., Pittman, A., Hodge, J.J.L., and Jepson, J.E.C. (2021). Impaired Pre-Motor Circuit Activity and Movement in a Drosophila Model of KCNMA1-Linked Dyskinesia. Movement Disorders 36, 1158–1169. 10.1002/mds.28479.

Lee, D., and O’Dowd, D.K. (1999). Fast excitatory synaptic transmission mediated by nicotinic acetylcholine receptors in Drosophila neurons. Journal of Neuroscience 19, 5311–5321.

Littleton, J.T., Stern, M., Perin, M., and Bellen, H.J. (1994). Calcium-dependence of neurotransmitter release and rate of spontaneous vesicle fusions are altered in Drosophila synaptotagmin mutants. Proceedings of the National Academy of Sciences of the United States of America 91, 10888–10892. 10.1073/pnas.91.23.10888.

Littleton, J.T., Stern, M., Schulze, K., Perin, M., and Bellen, H.J. (1993). Mutational analysis of Drosophila synaptotagmin demonstrates its essential role in Ca(2+)-activated neurotransmitter release. Cell 74, 1125–1134. 10.1016/0092-8674(93)90733-7.

Marchuk, D.A., Saulino, A.M., Tavakkol, R., Swaroop, M., Wallace, M.R., Andersen, L.B., Mitchell, A.L., Gutmann, D.H., Boguski, M., and Collins, F.S. (1991). cDNA cloning of the type-1 neurofibromatosis gene - complete sequence of the NF1 gene-product Genomics 11, 931–940. 10.1016/0888-7543(91)90017-9.

Mauthner, S.E., Hwang, R.Y., Lewis, A.H., Xiao, Q., Tsubouchi, A., Wang, Y., Honjo, K., Skene, J.H.P., Grandl, J., and Tracey, W.D. (2014). Balboa Binds to Pickpocket In Vivo and Is Required for Mechanical Nociception in Drosophila Larvae. Current Biology 24. 10.1016/j.cub.2014.10.038.

Melom, J.E., Akbergenova, Y., Gavornik, J.P., and Littleton, J.T. (2013). Spontaneous and Evoked Release Are Independently Regulated at Individual Active Zones. Journal of Neuroscience 33, 17253–17263. 10.1523/jneurosci.3334-13.2013.

Miles, J.H., Takahashi, T.N., Bagby, S., Sahota, P.K., Vaslow, D.F., Wang, C.H., Hillman, R.E., and Farmer, J.E. (2005). Essential versus complex autism: Definition of fundamental prognostic subtypes. American Journal of Medical Genetics Part A 135A, 171–180. 10.1002/ajmg.a.30590.

Molosh, A.I., Johnson, P.L., Spence, J.P., Arendt, D., Federici, L.M., Bernabe, C., Janasik, S.P., Segu, Z.M., Khanna, R., Goswami, C., et al. (2014). Social learning and amygdala disruptions in Nf1 mice are rescued by blocking p21-activated kinase. Nature Neuroscience 17, 1583–1590. 10.1038/nn.3822.

Morris, S.M., Acosta, M.T., Garg, S., Green, J., Huson, S., Legius, E., North, K.N., Payne, J.M., Plasschaert, E., Frazier, T.W., et al. (2016). Disease Burden and Symptom Structure of Autism in Neurofibromatosis Type 1 A Study of the International NF1-ASD Consortium Team (INFACT). Jama Psychiatry 73, 1276–1284. 10.1001/jamapsychiatry.2016.2600.

Mosca, T.J., Carrillo, R.A., White, B.H., and Keshishian, H. (2005). Dissection of synaptic excitability phenotypes by using a dominant-negative Shaker K+ channel subunit. Proceedings of the National Academy of Sciences of the United States of America 102, 3477–3482. 10.1073/pnas.0406164102.

Moscato, E.H., Dubowy, C., Walker, J.A., and Kayser, M.S. (2020). Social Behavioral Deficits with Loss of Neurofibromin Emerge from Peripheral Chemosensory Neuron Dysfunction. Cell Reports 32, 107856. 10.1016/j.celrep.2020.107856.

Nelson, S.B., and Valakh, V. (2015). Excitatory/Inhibitory Balance and Circuit Homeostasis in Autism Spectrum Disorders. Neuron 87, 684–698. 10.1016/j.neuron.2015.07.033.

Omrani, A., van der Vaart, T., Mientjes, E., van Woerden, G.M., Hojjati, M.R., Li, K.W., Gutmann, D.H., Levelt, C.N., Smit, A.B., Silva, A.J., et al. (2015). HCN channels are a novel therapeutic target for cognitive dysfunction in Neurofibromatosis type 1. Molecular Psychiatry 20, 1311–1321. 10.1038/mp.2015.48.

Parikshak, N.N., Luo, R., Zhang, A., Won, H., Lowe, J.K., Chandran, V., Horvath, S., and Geschwind, D.H. (2013). Integrative Functional Genomic Analyses Implicate Specific Molecular Pathways and Circuits in Autism. Cell 155, 1008–1021. 10.1016/j.cell.2013.10.031.

Powers, A.S., Grizzaffi, J., Ribchester, R., and Lnenicka, G.A. (2016). Regulation of quantal currents determines synaptic strength at neuromuscular synapses in larval Drosophila. Pflugers Archiv-European Journal of Physiology 468, 2031–2040. 10.1007/s00424-016-1893-7.

Ratner, N., and Miller, S.J. (2015). A RASopathy gene commonly mutated in cancer: the neurofibromatosis type 1 tumour suppressor. Nature Reviews Cancer 15, 290–301. 10.1038/nrc3911.

Ribeiro, M.J., Violante, I.R., Bernardino, I., Edden, R.A.B., and Castelo-Branco, M. (2015). Abnormal relationship between GABA, neurophysiology and impulsive behavior in neurofibromatosis type 1. Cortex 64, 194–208. 10.1016/j.cortex.2014.10.019.

Robertson, C.E., and Baron-Cohen, S. (2017). Sensory perception in autism. Nature Reviews Neuroscience 18, 671–684. 10.1038/nrn.2017.112.

Rubenstein, J.L.R., and Merzenich, M.M. (2003). Model of autism: increased ratio of excitation/inhibition in key neural systems. Genes Brain and Behavior 2, 255–267. 10.1034/j.1601-183X.2003.00037.x.

Russo, A., and DiAntonio, A. (2019). Wnd/DLK Is a Critical Target of FMRP Responsible for Neurodevelopmental and Behavior Defects in the Drosophila Model of Fragile X Syndrome. Cell Reports 28, 2581-+. 10.1016/j.celrep.2019.08.001.

Sanchez-Ortiz, E., Cho, W., Nazarenko, I., Mo, W., Chen, J., and Parada, L.F. (2014). NF1 regulation of RAS/ERK signaling is required for appropriate granule neuron progenitor expansion and migration in cerebellar development. Genes & Development 28, 2407–2420. 10.1101/gad.246603.114.

Shih, Y.T., Huang, T.N., Hu, H.T., Yen, T.L., and Hsueh, Y.P. (2020). Vcp Overexpression and Leucine Supplementation Increase Protein Synthesis and Improve Fear Memory and Social Interaction of Nf1 Mutant Mice. Cell Reports 31, 107835. 10.1016/j.celrep.2020.107835.

Shilyansky, C., Karlsgodt, K.H., Cummings, D.M., Sidiropoulou, K., Hardt, M., James, A.S., Ehninger, D., Bearden, C.E., Poirazi, P., Jentsch, J.D., et al. (2010). Neurofibromin regulates corticostriatal inhibitory networks during working memory performance. Proceedings of the National Academy of Sciences of the United States of America 107, 13141–13146. 10.1073/pnas.1004829107.

Shofty, B., Bergmann, E., Zur, G., Asleh, J., Bosak, N., Kavushansky, A., Castellanos, F.X., Ben-Sira, L., Packer, R.J., Vezina, G.L., et al. (2019). Autism-associated Nf1 deficiency disrupts corticocortical and corticostriatal functional connectivity in human and mouse. Neurobiology of Disease 130, 104479. 10.1016/j.nbd.2019.104479.

Stewart, B.A., Atwood, H.L., Renger, J.J., Wang, J., and Wu, C.F. (1994). Improved stability of Drosophila larval neuromuscular preparations in hemolymph-like physiological solutions. Journal of Comparative Physiology a-Sensory Neural and Behavioral Physiology 175, 179–191. 10.1007/bf00215114.

Sztainberg, Y., and Zoghbi, H.Y. (2016). Lessons learned from studying syndromic autism spectrum disorders. Nature Neuroscience 19, 1408–1417. 10.1038/nn.4420.

Takarae, Y., and Sweeney, J. (2017). Neural Hyperexcitability in Autism Spectrum Disorders. Brain Sciences 7, 129. 10.3390/brainsci7100129.

Tammimies, K., Marshall, C.R., Walker, S., Kaur, G., Thiruvahindrapuram, B., Lionel, A.C., Yuen, R.K.C., Uddin, M., Roberts, W., Weksberg, R., et al. (2015). Molecular Diagnostic Yield of Chromosomal Microarray Analysis and Whole-Exome Sequencing in Children With Autism Spectrum Disorder. Jama-Journal of the American Medical Association 314, 895–903. 10.1001/jama.2015.10078.

Tavassoli, T., Miller, L.J., Schoen, S.A., Nielsen, D.M., and Baron-Cohen, S. (2014). Sensory over-responsivity in adults with autism spectrum conditions. Autism 18, 428–432. 10.1177/1362361313477246.

The, I., Hannigan, G.E., Cowley, G.S., Reginald, S., Zhong, Y., Gusella, J.F., Hariharan, I.K., and Bernards, A. (1997). Rescue of a Drosophila NF1 mutant phenotype by protein kinase A. Science 276, 791–794. 10.1126/science.276.5313.791.

Tomchek, S.D., and Dunn, W. (2007). Sensory processing in children with and without autism: A comparative study using the short sensory profile. American Journal of Occupational Therapy 61, 190–200. 10.5014/ajot.61.2.190.

Tracey, W.D., Wilson, R.I., Laurent, G., and Benzer, S. (2003). painless, a Drosophila gene essential for nociception. Cell 113, 261–273. 10.1016/s0092-8674(03)00272-1.

Tsai, P.I., Wang, M.Y., Kao, H.H., Cheng, Y.J., Walker, J.A., Chen, R.H., and Chien, C.T. (2012). Neurofibromin Mediates FAK Signaling in Confining Synapse Growth at Drosophila Neuromuscular Junctions. Journal of Neuroscience 32, 16971–16981. 10.1523/jneurosci.1756-12.2012.

Valdez, C., Scroggs, R., Chassen, R., and Reiter, L.T. (2015). Variation in Dube3a expression affects neurotransmission at the Drosophila neuromuscular junction. Biology Open 4, 776–782. 10.1242/bio.20148045.

Violante, I.R., Patricio, M., Bernardino, I., Rebola, J., Abrunhosa, A.J., Ferreira, N., and Castelo-Branco, M. (2016). GABA deficiency in NF1 A multimodal C-11 -flumazenil and spectroscopy study. Neurology 87, 897–904. 10.1212/wnl.0000000000003044.

Violante, I.R., Ribeiro, M.J., Edden, R.A.E., Guimarares, P., Bernardino, I., Rebola, J., Cunha, G., Silva, E., and Castelo-Branco, M. (2013). GABA deficit in the visual cortex of patients with neurofibromatosis type 1: genotype-phenotype correlations and functional impact. Brain 136, 918–925. 10.1093/brain/aws368.

Vorstman, J.A.S., Spooren, W., Persico, A.M., Collier, D.A., Aigner, S., Jagasia, R., Glennon, J.C., and Buitelaar, J.K. (2014). Using genetic findings in autism for the development of new pharmaceutical compounds. Psychopharmacology 231, 1063–1078. 10.1007/s00213-013-3334-z.

Walcott, K.C.E., Mauthner, S.E., Tsubouchi, A., Robertson, J., and Tracey, W.D. (2018). The Drosophila Small Conductance Calcium-Activated Potassium Channel Negatively Regulates Nociception. Cell Reports 24, 3125-+. 10.1016/j.celrep.2018.08.070.

Walker, J.A., Gouzi, J.Y., Long, J.B., Huang, S.D., Maher, R.C., Xia, H.J., Khalil, K., Ray, A., Van Vactor, D., Bernards, R., and Bernards, A. (2013). Genetic and Functional Studies Implicate Synaptic Overgrowth and Ring Gland cAMP/PKA Signaling Defects in the Drosophila melanogaster Neurofibromatosis-1 Growth Deficiency. Plos Genetics 9, e1003958. 10.1371/journal.pgen.1003958.

Walker, J.A., Tchoudakova, A.V., McKenney, P.T., Brill, S., Wu, D.Y., Cowley, G.S., Hariharan, I.K., and Bernards, A. (2006). Reduced growth of Drosophila neurofibromatosis 1 mutants reflects a non-cell-autonomous requirement for GTPase-activating protein activity in larval neurons. Genes & Development 20, 3311–3323. 10.1101/gad.1466806.

Wang, Y.Y., Brittain, J.M., Wilson, S.M., Hingtgen, C.M., and Khanna, R. (2010). Altered calcium currents and axonal growth in Nf1 haploinsufficient mice. Translational Neuroscience 1, 106–114. 10.2478/v10134-010-0025-8.

Williams, J.A., Su, H.S., Bernards, A., Field, J., and Sehgal, A. (2001). A circadian output in Drosophila mediated by neurofibromatosis-1 and Ras/MAPK. Science 293, 2251–2256. 10.1126/science.1063097.

Willsey, A.J., Sanders, S.J., Li, M.F., Dong, S., Tebbenkamp, A.T., Muhle, R.A., Reilly, S.K., Lin, L., Fertuzinhos, S., Miller, J.A., et al. (2013). Coexpression Networks Implicate Human Midfetal Deep Cortical Projection Neurons in the Pathogenesis of Autism. Cell 155, 997–1007. 10.1016/j.cell.2013.10.020.

Yenkoyan, K., Grigoryan, A., Fereshetyan, K., and Yepremyan, D. (2017). Advances in understanding the pathophysiology of autism spectrum disorders. Behavioural Brain Research 331, 92–101. 10.1016/j.bbr.2017.04.038.

Zhang, Y., Bonnan, A., Bony, G., Ferezou, I., Pietropaolo, S., Ginger, M., Sans, N., Rossier, J., Oostra, B., Le Masson, G., and Frick, A. (2014). Dendritic channelopathies contribute to neocortical and sensory hyperexcitability in Fmr1-(/y) mice. Nature Neuroscience 17, 1701–1709. 10.1038/nn.3864.

Zoghbi, H.Y., and Bear, M.F. (2012). Synaptic Dysfunction in Neurodevelopmental Disorders Associated with Autism and Intellectual Disabilities. Cold Spring Harbor Perspectives in Biology 4, a009886. 10.1101/cshperspect.a009886.

