## Supplemental Information Figures and Legends for "Loss of *NF1* causes tactile hypersensitivity and impaired synaptic transmission in a *Drosophila* model of autism spectrum disorder"

**Video S1. Typical K33 response to mechanical stimulation.** Upon brief mechanical stimulation, K33 larvae typically do not display the stereotypic larval nocifensive response.

**Video S2. Typical** ***$NF1^{P1}$*** **response to mechanical stimulation.** Upon brief mechanical stimulation, *$NF1^{P1}$* larvae typically display the stereotypic larval nocifensive response.

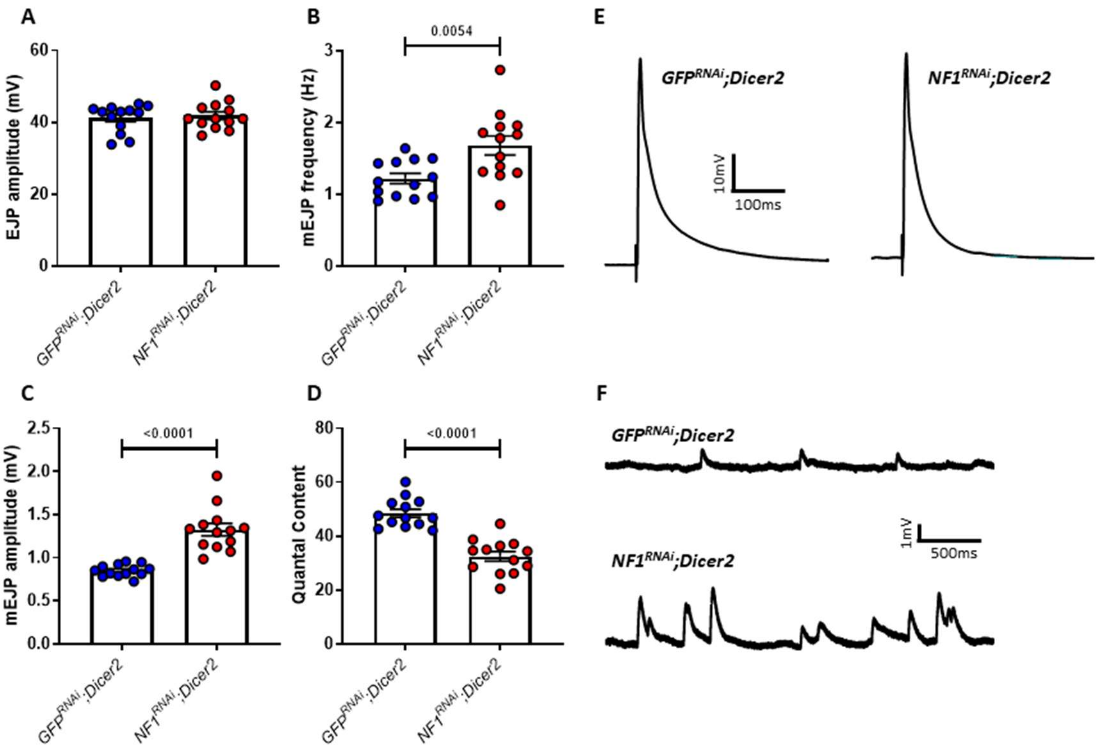


**Figure S1. Pan-neuronal knockdown of *NF1* via an alternative *UAS-NF1$^{RNAi}$* construct (VDRC #35077) mimics the mutant phenotype.** *A)* EJP amplitude is unaffected by pan-neuronal (*elav*-driven) *NF1$^{RNAi}$;Dicer2* expression (p=0.66). *B)* Knockdown of *NF1* increases mEJP frequency and *C)* mEJP amplitude, with a significant reduction in *D)* quantal content. *E-F)* Representative traces of EJPs and mEJPs, respectively. All data are presented as mean ± SEM. All statistical comparisons were made via unpaired, two-tailed student's *t*-test.

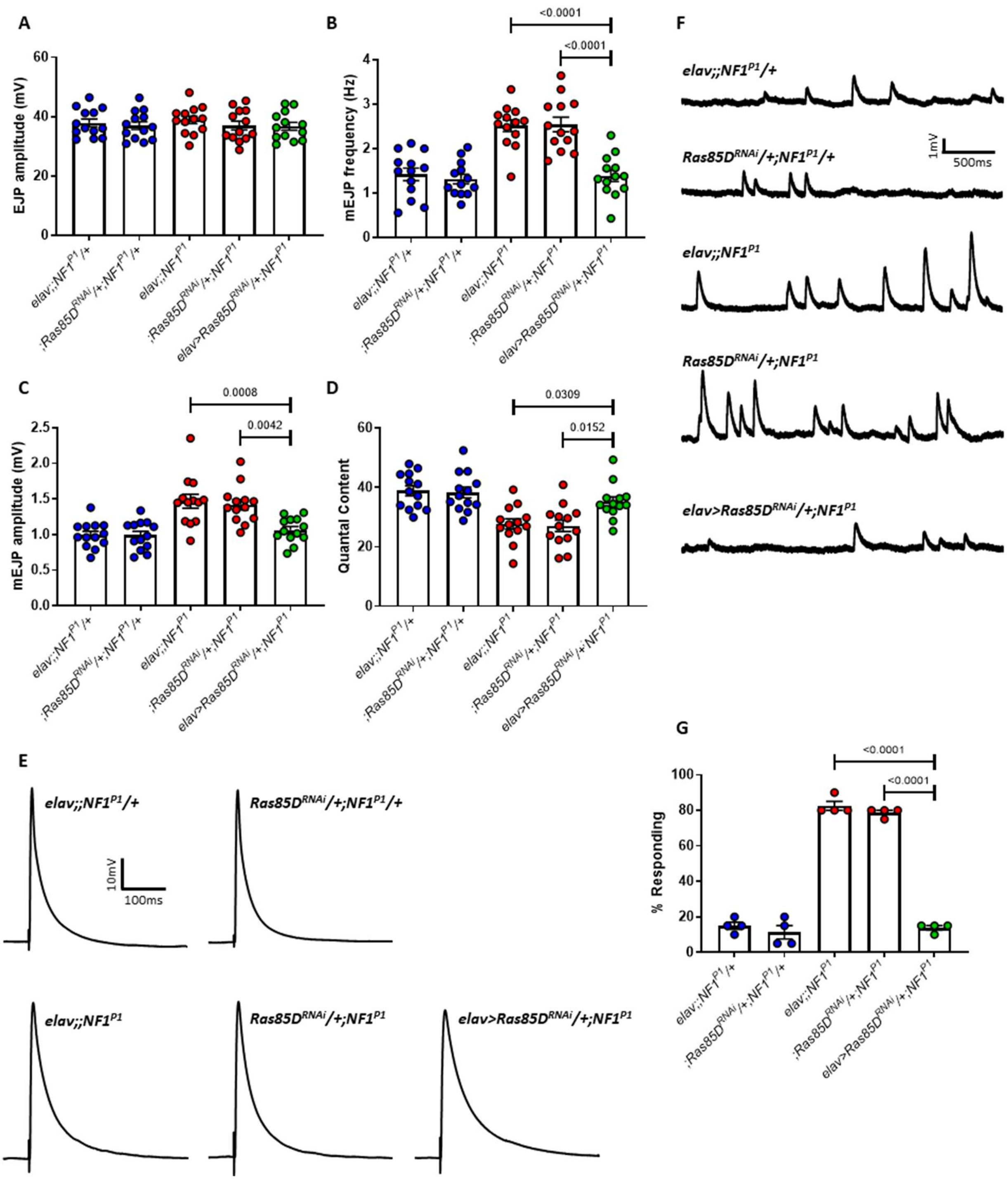


**Figure S2. Knockdown of *Ras85D* rescues synaptic transmission deficits and tactile hypersensitivity in *NF1*$^{P1}$ larvae.** *A)* EJP amplitude was not significantly different between any of the lines tested. *B)* mEJP frequency in *elav>Ras85D*$^{RNAi}$*/+;NF1*$^{P1}$ larvae (rescue line; green circles) is significantly reduced compared to both homozygous mutant lines (red circles). Expression of UAS-*Ras85D*$^{RNAi}$ also rescues *C)* increased mEJP amplitude and *D)* reduced quantal content. There were no significant differences between the rescue line and either heterozygous control (blue circles) for any parameter examined. Furthermore, in panels B-D, both heterozygous controls were significantly different to both homozygous mutant controls, and there were no significant differences between either of the heterozygous controls or either of the homozygous mutant controls, respectively. *E-F)* Representative traces of EJPs and mEJPs, respectively, for each of the lines tested in A-D. G) Pan-neuronal expression of *UAS-Ras85D*$^{RNAi}$ is sufficient to rescue tactile hypersensitivity in *NF1*$^{P1}$ larvae. All data are presented as mean ± SEM. All statistical comparisons were made via a one-way ANOVA followed by Tukey's post-hoc test.

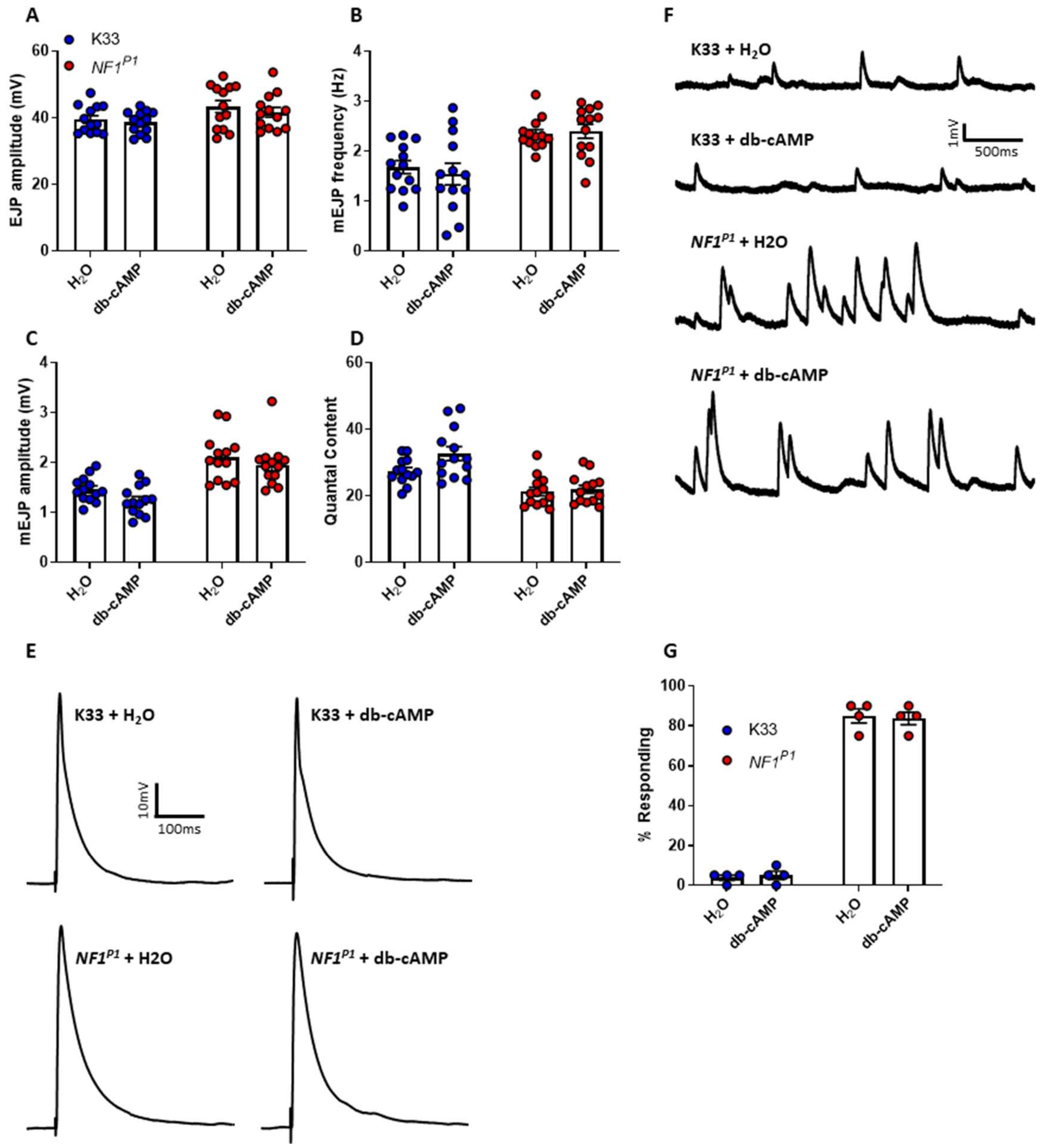


**Figure S3. Raising *NF1*$^{P1}$ larvae on db-cAMP does not rescue impaired synaptic transmission or tactile hypersensitivity**. *In* *NF1*$^{P1}$ larvae, 10 µM db-cAMP administered throughout development does not significantly affect *A)* EJP amplitude ($p=0.81$), *B)* mEJP frequency ($p=0.99$), *C)* mEJP amplitude ($p=0.75$), or *D)* quantal content ($p=0.99$). The apparent increase in quantal in K33 larvae following db-cAMP treatment is non-significant ($p=0.070$). In panels B-D, $H_2O$-treated *NF1*$^{P1}$larvae are significantly different to $H_2O$-treated K33 larvae. *E-F)* Representative traces of EJPs and mEJPs, respectively. *G)* Likewise, there is no effect of db-cAMP treatment on tactile hypersensitivity in *NF1*$^{P1}$ larvae ($p=0.99$). All data are presented as mean ± SEM. All statistical comparisons were made via two-way ANOVA followed by Tukey's multiple comparisons test, in which each genotype + treatment was compared to all others.
